# Editing cis-regulatory elements towards generating rice stomatal morphological variation for adaptation to broad and dynamic environments

**DOI:** 10.1101/2023.11.29.569268

**Authors:** Nicholas G. Karavolias, Dhruv Patel-Tupper, Ana Gallegos Cruz, Lilian Litvak, Samantha E. Lieberman, Michelle Tjahjadi, Krishna K. Niyogi, Myeong-Je Cho, Brian J. Staskawicz

**Affiliations:** Innovative Genomics Institute, Berkeley, CA 94720; Plant and Microbial Biology Department, UC Berkeley, Berkeley, CA 94720; Howard Hughes Medical Institute, University of California, Berkeley, CA 94720, USA; Molecular Biophysics and Integrated Bioimaging Division, Lawrence Berkeley National Laboratory, Berkeley, CA 94720, USA

## Abstract

Cis-regulatory element editing can generate quantitative trait variation while mitigating against extreme phenotypes and harmful pleiotropy associated with coding sequence mutations. Here, we applied a multiplexed guide RNA design approach, informed by bioinformatic datasets, to generate genotypic variation in the promoter of *OsSTOMAGEN,* a positive regulator of stomatal density in rice. Engineered genotypic variation corresponded to broad and continuous variation in stomatal density, ranging from 70% to 120% of wild-type stomatal density. This near-isogenic panel of stomatal variants was leveraged in physiological assays to establish discrete relationships between stomatal morphological variation and stomatal conductance, carbon assimilation, and intrinsic water use efficiency in steady-state and fluctuating light conditions. Additionally, promoter alleles were subjected to vegetative drought regimes to assay the effects of the edited alleles on developmental response to drought. Notably, the capacity for drought-responsive stomatal density reprogramming in *stomagen* and two cis-regulatory edited alleles was reduced. Collectively our data demonstrate that cis-regulatory element editing can generate near-isogenic trait variation that can be leveraged for establishing relationships between anatomy, physiology, and crop improvement along diverse environmental clines.

## Introduction

In a global food system threatened by the effects of climate change on water availability, stomata have emerged as a central target of engineering due to their seminal role in water loss and photosynthesis^1^. Efforts to adapt germplasm to variable environments often include modifications to stomatal characteristics including density and kinetics^2–5^. Generating stomatal variants adapted to diverse environments remains a highly sought-after goal.

The influence of stomatal density on other stomatal morphological traits, gas exchange, intrinsic water use efficiency (iWUE), and drought tolerance has been shown to be significant in a broad array of species^2,3,6–10^. In rice specifically, reductions in stomatal density have been associated with increases in iWUE, water conservation, and drought adaptation^2,3,8,11^. Despite promising abiotic stress tolerance phenotypes, rice with reduced stomatal densities sometimes exhibit concomitant reductions of stomatal conductance and carbon assimilation which may lower overall productivity^2,8,11,12^. Studies seeking to relate stomatal density with physiology have also been constrained by limited morphological variation and genetically heterogenous backgrounds that complicate the ability to parse the effects of anatomical variation from the effects of other genetic determinants ^2,3,7–9^. In order to effectively leverage stomatal density alterations for climate change resilience, greater insights into the physiological outcomes of these modifications are needed.

Gene editing in plant species spurred on by the introduction of CRISPR/Cas tools has been leveraged to generate variation in a breadth of climate-resilience traits, including stomata^13^. The vast majority of successes to date have been achieved through gene knockouts^13^. In some cases, a knockout-based approach can provide sufficient trait variation while minimizing deleterious pleiotropic effects^8,13^. This is especially true in cases where paralogs exist^8,14,15^. However, this approach is limited to a small pool of genes whose null phenotypes are not detrimental to overall plant fitness.

Editing *cis*-regulatory elements is an attractive, alternative, gene-editing approach to generate novel variation. Targeting mutations to regulatory regions can fine-tune expression of genes towards more subtle modifications to plant phenotypes with fewer pleiotropic effects relative to coding sequence mutations^15–17^. Beyond the potential for crop improvement, *cis*-regulatory element editing can help provide insights into the complex and obscure process of transcriptional regulation.

To generate variation in stomatal traits, we designed guide RNAs targeted to the promoter of STOMAGEN in rice. *OsSTOMAGEN*, is a positive regulator of stomatal development, whose knockout produces severe reductions in stomatal density ^8,18^. Promoter editing was used to mitigate this extreme knockout phenotype and produce quantitative variation in stomatal density. Rice production landscapes encompass a broad range of environmental conditions with radically variable water availability^19,20^. Cultivated varieties of rice display a range of stomatal densities and morphologies corresponding to the environment where they are produced^3,21^. The variation generated in this study was used to interrogate the structure-function relationships of stomatal density and physiology and the capacity of *cis*-regulatory mutagenesis to fine-tune stomatal density for adaptation to broad rice production systems.

Our work applies a multiplexed guide design approach for producing quantitative trait variation in rice stomatal density. The transcriptional and phenotypic outcomes of cis-regulatory editing of *OsSTOMAGEN* are measured. The quantitative trait variation established is leveraged to investigate the relationship of stomatal density to gas exchange in a near-isogenic population.

Finally, we consider stomatal density reprogramming under vegetative drought stress to assess the impacts of *cis*-regulatory element editing on environmental response. We apply these findings towards a framework to understand how approaches in promoter editing can be used to established quantitative trait variation for broad and dynamic environments.

## Methods

### Plant growth conditions

Rice cultivar Nipponbare (*O. sativ*a ssp. japonica) seeds were germinated and grown for 8 days in a petri dish with 20 mL of water in a Conviron growth chamber at 28**°**C for day-length periods of 16 hours in 100 μmol photons m^-2^ s^-1^ of light and 80% relative humidity. Seedlings were transferred to a soil mixture comprised of equal parts turface (https://www.turface.com/products/infield-conditioners/mvp) and sunshine mix #4 (http://www.sungro.com/professional-products/fafard/). Germinated seedlings used for stomatal phenotyping and growth chamber physiological assays were transferred to 10 cm, 0.75 L McConkey tech square pots and placed in growth chambers at 28**°**C for day-length periods of 16 hours in 400 μmol photons m^-2^ s^-1^ of light and 80% relative humidity.

Plants designated for greenhouse measurements were placed in setpoints of 27**°**C day/22°C night at ambient light conditions with day lengths of 12 hours in 15.2 cm, 1.8 L pots. All plants were fertilized with 125 mL of 1% w/v iron solution one-week post-transplant. 1 L of 5% w/v JR Peter’s Blue 20-20-20 fertilizer (https://www.jrpeters.com/) was added to each flat at 3- and 11-weeks post-germination.

### Identifying putative transcription factor binding sites, conserved non-coding sequences, regions of open chromatin, and H3K27ac DNA interaction regions

Transcription factor binding sites were identified using the total available transcription factor binding site motifs in JASPAR Plant Core^22^. The region 2 kb upstream of the translation start site was queried for the existence of transcription factor binding sites using a 99% score threshold.

Conserved non-coding sequences within the Poaceae family were identified using mVista^23^. The sequences of the 2 kb region upstream of the translation start site of the homolog of *OsSTOMAGEN* from *Brachypodium distachyon*, *Hordeum vulgare*, *Setaria italica,* and *Zea mays* was retrieved from Phytozome. A duplication of *OsSTOMAGEN* in Poaceae required the use of the gene tree generated by Phytozome to identify orthologs of *STOMAGEN* within each species assayed. All *STOMAGEN* copies also had the highest percent identity compared to *OsSTOMAGEN* relative to other paralogs of *STOMAGEN* in species assayed. ATAC-seq and ChIP-seq data of *OsSTOMAGEN* were extracted from the publicly available RiceENCODE database (http://glab.hzau.edu.cn/RiceENCODE/pages/Browser.html)^24^.

### Generation of edited lines

Forward and reverse strand guide sequences with relevant sticky ends amenable for Golden Gate cloning were ordered from Integrated DNA Technology (IDTdna.com). Guide sequences are listed in Table S1. Guides are arranged in order from most distal to proximal of the Os*STOMAGEN* transcription start site. Equal volumes of 10 mM primers were annealed at room temperature. Golden Gate cloning was used to insert all eight guides simultaneously into the entry clone containing the tracrRNA and U3 promoter. LR clonase reactions were used to insert the entry clone into destination vectors for biolistic transformation and *Agrobacterium-*mediated transformation. The guide for targeting the coding sequence of *OsSTOMAGEN* was selected to minimize off-targeting, using CRISPR-P 2.0 (http://crispr.hzau.edu.cn/CRISPR2/).

### Plant material and culture of explants

Mature seeds of rice (*O. sativa* L. japonica cv. Kitaake) were de-hulled, and surface-sterilized for 20 min in 20% (v/v) commercial bleach (5.25% sodium hypochlorite) and 1% of Tween 20. Three washes in sterile water were used to remove residual bleach from seeds. De-hulled seeds were placed on callus induction medium (CIM) medium [N6 salts and vitamins^25^, 30 g/L maltose, 0.1 g/L myo-inositol, 0.3 g/L casein enzymatic hydrolysate, 0.5 g/L L-proline, 0.5 g/L L-glutamine, 2.5 mg/L 2,4-D, 0.2 mg/L BAP, 5 mM CuSO4, 3.5 g/L Phytagel, pH 5.8] and incubated in the dark at 28°C to initiate callus induction. Six- to 8-week-old embryogenic calli were used as targets for transformation.

### *Agrobacterium*-mediated transformation

Embryogenic calli were dried for 30 min prior to incubation with an *Agrobacterium tumefaciens* EHA105 suspension (OD_600_ = 0.1) carrying the binary vector for editing rice STOMAGEN promoter. After a 30 min incubation, the *Agrobacterium* suspension was removed. Calli were then placed on sterile filter paper, transferred to co-cultivation medium [N6 salts and vitamins, 30 g/L maltose, 10 g/L glucose, 0.1 g/L myo-inositol, 0.3 g/L casein enzymatic hydrolysate, 0.5 g/L L-proline, 0.5 g/L L-glutamine, 2 mg/L 2,4-D, 0.5 mg/L thiamine, 100 mM acetosyringone, 3.5 g/L Phytagel, pH 5.2] and incubated in the dark at 21°C for 3 days. After co-cultivation, calli were transferred to resting medium [N6 salts and vitamins, 30 g/L maltose, 0.1 g/L myo-inositol, 0.3 g/L casein enzymatic hydrolysate, 0.5 g/L L-proline, 0.5 g/L L-glutamine, 2 mg/L 2,4-D, 0.5 mg/L thiamine, 100 mg/L timentin, 3.5 g/L Phytagel, pH 5.8] and incubated in the dark at 28°C for 7 days. Calli were then transferred to selection medium [CIM plus 250 mg/L cefotaxime and 50 mg/L hygromycin B] and allowed to proliferate in the dark at 28°C for 14 days. Well-proliferating tissues were transferred to CIM containing 75 mg/l hygromycin B. The remaining tissues were subcultured at 3 to 4 week intervals on fresh selection medium. When a sufficient amount (about 1.5 cm in diameter) of the putatively transformed tissues was obtained, they were transferred to regeneration medium [MS salts and vitamins^26^, 30 g/L sucrose, 30 g/L sorbitol, 0.5 mg/L NAA, 1 mg/L BAP, 150 mg/L cefotaxime] containing 40 mg/L hygromycin B and incubated at 26°°C, 16-h light, 90 μmol photons m^-2^ s^-1^. When regenerated plantlets reached at least 1 cm in height, they were transferred to rooting medium [MS salts and vitamins, 20 g/L sucrose, 1 g/L myo-inositol, 150 mg/L cefotaxime] containing 20 mg/L hygromycin B and incubated at 26^°o^C under conditions of 16-h light (150 μmol photons m^-2^ s^-1^) and 8-h dark until roots were established and leaves touched the Phytatray^TM^ II lid (Sigma-Aldrich, St. Louis, MO, USA). Plantlets were then transferred to soil.

### Validation of edits

T_0_ plants targeted for edits in Os*STOMAGEN* promoter were evaluated using PCR to amplify the entire 2 kb region upstream of the translation start site. To account for the possibility of heterozygous promoter mutations PCR products were subcloned into Zero Blunt™ TOPO™ (Thermo Fisher, Waltham, MA). Fifteen *E. coli* transformants were miniprepped and sequenced using theprimers listed in Table 1. Seeds from T_o_ plants with heterozygous promoter mutations were genotyped using the same method in the T_1_ generation to isolate homozygous promoter mutations. Azygous T_2_ plants were used for experimental data collection to minimize somaclonal variation, which may have accumulated during tissue culture^27,28^.

Edits in the coding sequence of *OsSTOMAGEN* were evaluated using PCR to amplify the region of interest (Table S2). PCR products were Sanger sequenced. Sequence data was analyzed using the Synthego ICE tool (https://ice.synthego.com/#/) to detect alleles present^29^. Only lines with homozygous frame-shift mutations were retained for downstream experiments

### Phenotyping stomatal density and size

Stomatal densities were recorded from epidermal impressions of leaves using nail polish peels^30^. Impressions were made on the widest section of 32-day-old fully expanded leaves of each biological replicate. Images were taken using a Leica DM5000 B epifluorescence microscope at 10x magnification. Three images were collected per stomatal impression and density per image was averaged. The number of stomata in a single stomatal band were counted and the area of each band was measured^31^. Stomatal densities were calculated by dividing stomatal counts by stomal band area (mm^-2^). Stomatal densities of fifth leaf abaxial tissues were assayed for each allele.

Stomatal density of leaves that developed during vegetative drought was measured using the same methods described above. The vegetative drought was applied by removing all water from the flats. Emerging leaves from the three largest tillers were marked four days after drought initiation. Drought was applied for a total of 7 days prior to rehydrating. Marked leaves were allowed to expand fully prior to making epidermal impressions.

Epidermal peels of 40-day old plants were produced using a razor blade on the adaxial leaf to remove tissues above the abaxial epidermal layer. Images of individual stomata at 100x magnification were captured. Guard cell length was measured using ImageJ. 30 individual stomata collected from a total of five biological replicates of each genotype were measured.

### Quantifying OsSTOMAGEN transcript abundance

Total RNA was extracted from plants with the Qiagen Total RNAeasy Plant Kit at three developmental stages: five days after germination, from the basal 2.5 cm of the youngest developing leaf on 32-day old plants, and 2.5 cm from the leaf tip of the flag leaf of the primary tiller from 64-day old plants for each promoter allele and wild type. For comparisons of transcript abundance in vegetative drought, total RNA was also extracted from the basal 2.5 cm of the youngest developing leaves of 32-day old plants after a 5-day vegetative drought stress was imposed. Flats of plants to be droughted were drained of all water five days prior to a sampling RNA was reverse transcribed using the QuantiTect^TM^ reverse transcription kit to generate first-strand cDNA. Quantitative reverse transcription PCR was performed using FAST SYBR on Applied Biosystem’s QuantStudio 3 thermocycler. Relative expression levels were calculated by normalizing to the average of rice housekeeping genes, *UBQ5* (LOC_Os01g22490) and *eEF-1A* (LOC_Os03g08020). ^32^. Primers used for qPCR listed in Supporting Table 1. Relative log fold expression was calculated using the 2^-ΔΔCT^method using “Do my qpcr” web tool ^33^. For flag leaves, seedlings, and developing leaves, all comparisons are made to wild type. For comparisons of well-watered to vegetative drought developing leaves all expression is relative to wild type well-watered. For genotype specific comparisons across tissue types, comparisons were made to flag leaves.

### Photosynthesis and stomatal conductance assays

Physiological assays in Figures 2 and 3 were conducted on the fully expanded leaf 5 of 32-day old plants. Data for Figures 2 and 3 was captured using an infrared gas analyzer (LI6800XT, LI-COR, Lincoln, NE, USA) with chamber conditions set to: light intensity 1500 μmol photons m^-2^ s^-1^ (90% red light, 10% blue light); leaf temperature 25°C; flow rate 500 μmol s^-1^; relative humidity 50%; and CO_2_ concentration of sample 400 ppm. Each biological replicate was allowed to acclimate for 15 minutes prior to data collection. Leaves from which gas exchange data was captured were marked and subsequently phenotyped for stomatal density.

Measurements of gas exchange on vegetatively droughted plants were conducted using the methods described above. Drought was applied by draining flats of plants entirely. Gas exchange measurements were taken on the fifth day of drought.

### Deriving stomatal limitation, J_max_, and V_cmax_ from A/Ci curves

Curves of carbon assimilation rates relative to substomatal CO_2_ concentrations were produced using the Dynamic Assimilation Technique ^TM^ (https://www.licor.com/env/support/LI-6800/topics/dynamic-assimilation-technique.html) using a LI6800 infrared gas analyzer (LI-COR, Lincoln, NE, USA) ^34^. Chamber parameters were set to the following parameters: leaf temperature 25°C; flow rate 500 μmol s^-1^; relative humidity 50%; light intensity 1500 μmol photons m^-2^ s^-1^; fan speed 10,000 rpm. CO concentration was varied from 10 to 1800 ppm with a ramp rate of 150 ppm per minute. Data was logged every four seconds. Data from a total of four biological replicates per genotype were collected.

A curve relating carbon assimilation to substomatal carbon dioxide concentration (C_i_) was fit to each biological replicate using the “Plantecophys” package in R Studio for all values of carbon assimilation greater than zero^35^. A C_i_ threshold of 700 ppm was used for modeling the curves relevant to the regions where stomatal limitation occurs.

Stomatal limitations were calculated using the differential method^36^. Briefly, the carbon assimilation rate predicted by the model when Ci equals the ambient carbon dioxide concentration set at 370 ppm was subtracted from the carbon assimilation rate predicted by the model when Ci=370 ppm. This difference was divided by the carbon assimilation rate predicted by the model when Ci=370 ppm and multiplied by 100 to represent the percentage of stomatal limitation. Stomatal limitations were calculated independently for each biological replicate.

A-Ci curves were also fit for the entire dynamic range of C_i_ values in order to estimate J_max_ and V_cmax_. All default parameters were used to model A-Ci curves.

### Fluctuating light assay

Plants were dark acclimated for one hour prior to light induction assay. Plants were then subjected to the following conditions leaf temperature 25°C; flow rate 500 μmol s^-1^; relative humidity 50%; and CO_2_ concentration of sample 400 ppm. The light intensity was initially set to 0 μmol photons m^-2^ s^-1^ for 3 minutes during which Fv/Fm recordings were made. Only plants with Fv/Fm greater than 0.78 were used in this assay. After 3 minutes, the light intensity was increased to 100 μmol photons m^-2^ s^-1^ for minutes after which high light was induced. Plants were kept at 1500 μmol photons m^-2^ s^-1^ (90% red light, 10% blue light) for 78 minutes prior to a return to 100 μmol photons m^-2^ s^-1^ for 15 more minutes. Data was logged every 10 seconds.

A curve fitting each data point to its adjacent point was produced and the area under this curve was calculated for stomatal conductance and for carbon assimilation. The integrated carbon assimilation value was divided by the integrated stomatal conductance value to generate a measurement of intrinsic water use efficiency in this dynamic range. A one phase decay model was fitted to the stomatal conductance data restricted to the region of the curve immediately after the light to dark transition. The rate constant of this curve was calculated and used to estimate stomatal closure rates.

### Graphs and statistics

Plots were generated using Adobe Illustrator 24.1. and Prism 9.5. Statistics were computed in R studio. Comparisons of multiple groups were conducted using ANOVA and Tukey’s honest significant difference (HSD) post-hoc tests.

## Results

### Bioinformatic analyses inform promoter regions for editing

Overlaying putative transcription factor binding sites, conserved non-coding sequences (CNS), ATAC-seq and ChIP-seq data indicated genomic signatures to target by cis-regulatory mutagenesis (Supplemental Figure 1). The greatest conservation among non-coding sequences was observed in *Brachypodium distachyon* relative to other family members. Transcription factor binding sites were the least harmonious across the assessed datasets.

The eight guide RNAs selected for targeting the 5’ upstream region of *OsSTOMAGEN* are indicated by blue triangles in Supplemental Figure 1e. All selected guides correspond to at least one signature detected in the bioinformatic analysis, prioritizing regions where ATAC-seq, ChiIP-seq, and CNSs peaks coincided. Alleles generated (Figure 1a) represent editing outcomes consistent with simultaneous and asynchronous activity of multiple guides. A single guide construct was used to generate coding sequence mutations in the first exon *OsSTOMAGEN* (Supplemental Figure 2).

**Figure 1.**
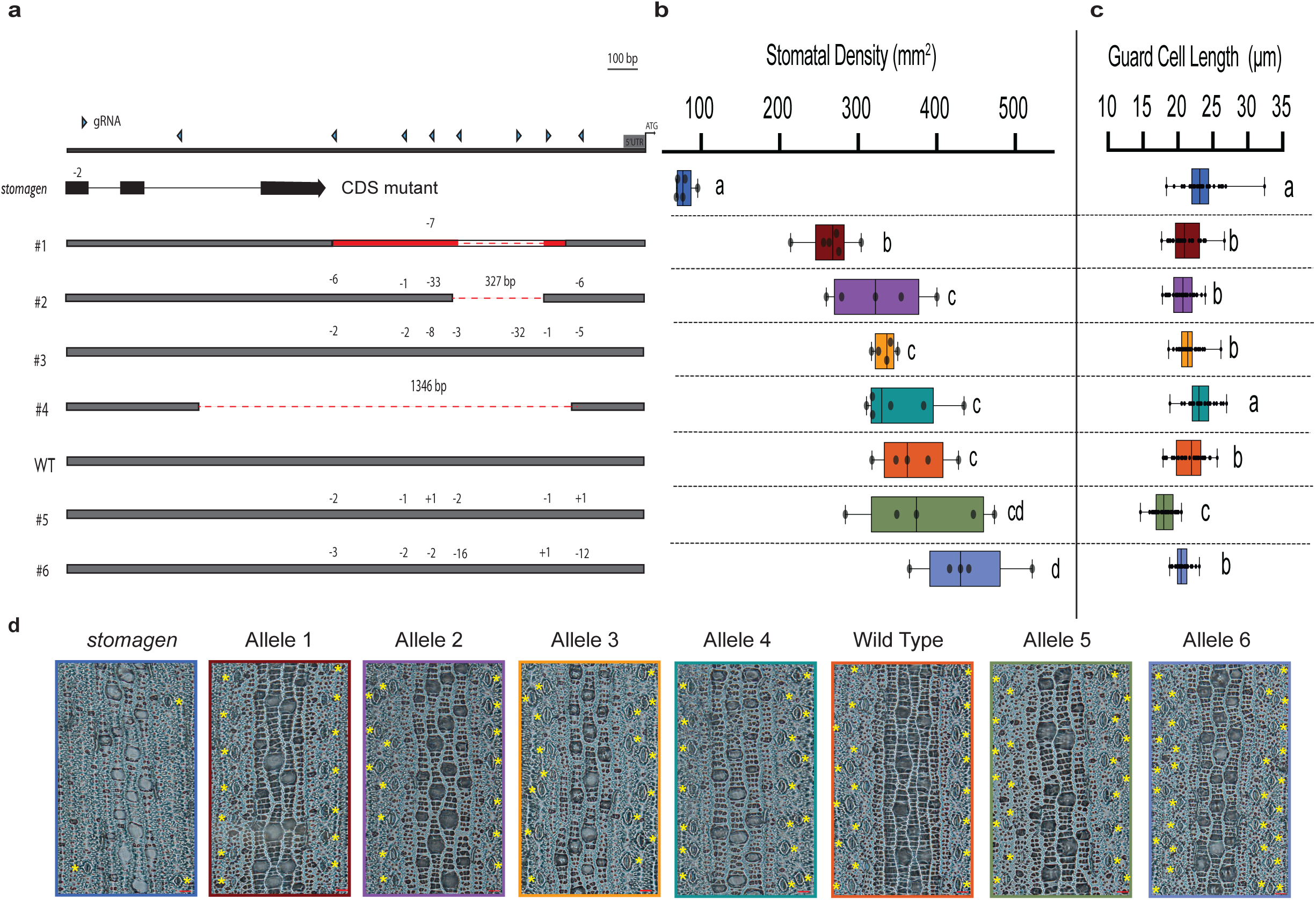
Stomatal density and variation in size across gene-edited *OsSTOMAGEN* promoter alleles. (a) The genotype of each promoter allele and the *stomagen* CDS mutant. Red shaded regions indicate inversion. The size of each indel is listed above cut site. (b) Box-and-whisker plot of the stomatal density of each allele assayed (c) Box-and-whisker plots of guard cell length of each allele assayed. (d) Representative images of each genotype assayed. Yellow asterisks are placed adjacent to each stoma. Red scale bars represent 20 μm. In the box-and-whisker plots, the center horizontal indicates the median, upper and lower edges of the box are the upper and lower quartiles and whiskers extend to the maximum and minimum values within 1.5 interquartile ranges. Each dot represents a biological replicate. Letters indicate a significant difference between means (P<0.05, one-way ANOVA Tukey HSD post-hoc test). Barplots mean is represented with error bars showing standard error of the mean. Asterisks represent a significant difference in expression relative to wild type (p<0.05, one-way ANOVA Tukey HSD post-hoc test).

### *OsSTOMAGEN* promoter alleles alter stomatal density and morphology

The phenotypic implications of *OsSTOMAGEN* promoter and coding sequence mutations were first assessed by measuring stomatal density. The stomatal density of the *stomagen* knockout in *Oryza sativa* cv. Kitaake exhibited an 80% reduction of wild type stomatal density, consistent with knockout phenotypes observed in cv. Nipponbare and cv. IR64 ^8,18^ (Figure 1b, Supplemental Figure 3). The panel of promoter edited alleles exhibited a broad diversity of stomatal densities including four lines with densities lower than— and two lines with densities greater than — wild type (Figure 1a, 1b Supplemental Figure 3). The ranked order of stomatal density alleles remained the same in greenhouse and growth chamber conditions (Figure 1b, Supplemental Figure 3).

Beyond density, guard cell length was measured as a proxy of stomatal size. Variation in stomatal size was observed in the panel of *OsSTOMAGEN* alleles assayed (Figure 1c). An overall negative correlation between stomatal density and size was present. Some deviations from a linear relationship suggest that stochasticity in guard cell length may be related to promoter mutations (Supplemental Figure 4). Broad quantitative variation in stomatal density and size was successfully established by coding sequence and promoter editing of *OsSTOMAGEN*.

### *OsSTOMAGEN* transcript abundance across multiple tissue types does not correlate with stomatal density phenotypes

The transcript abundance of *OsSTOMAGEN* across many tissue types was subsequently assessed in promoter alleles to determine the relationships between expression and phenotype. *OsSTOMAGEN* transcript abundance was measured in each promoter allele relative to wild type in flag leaves, seedlings, and developing leaves (Extended Data 1). In developing leaves where *OsSTOMAGEN* exhibits the greatest expression in wild type (Extended Data 1a), there is no obvious co-linearity of stomatal density and expression level. Similarly, in flag leaves and seedlings there is no obvious relationship between density and expression levels (Extended Data Fig 1b and 1c). The ratio of STOMAGEN expression levels within tissue types varied among promoter alleles as well (Extended Data 2).

### Near-isogenic panel reveals relationships between stomatal density and gas exchange physiology

The anatomical diversity of stomatal density warranted further assessments of physiology within a near-isogenic context. The relationships of stomatal density to carbon assimilation, stomatal conductance, iWUE, the quantum yield of photosystem II (ΦPSII), maximum rate of electron transport (J_max_) and maximum rate of carboxylation (V_cmax_) were established leveraging the diversity inherent to the panel (Figure 2). A strong, positive, linear association between stomatal density and steady-state carbon assimilation, stomatal conductance, and ΦPSII at a light intensity of 1500 μmol photons m^-2^ s^-1^ was observed (Figure 2a,b,d). Likewise, a strong, negative, linear association between stomatal density and intrinsic water-use efficiency was established (Figure 2c). Stomatal limitation, derived from A/C_i_ curves, in *stomagen* far exceeded other promoter alleles (Supplemental Figure 6). Limited variation in stomatal limitation was observed among promoter alleles. Measurements of J_max_ and V_cmax_ derived from A-Ci curves indicate some possible co-variation with stomatal density constrained by biological variation (Supplemental Figure 7).

**Figure 2.**
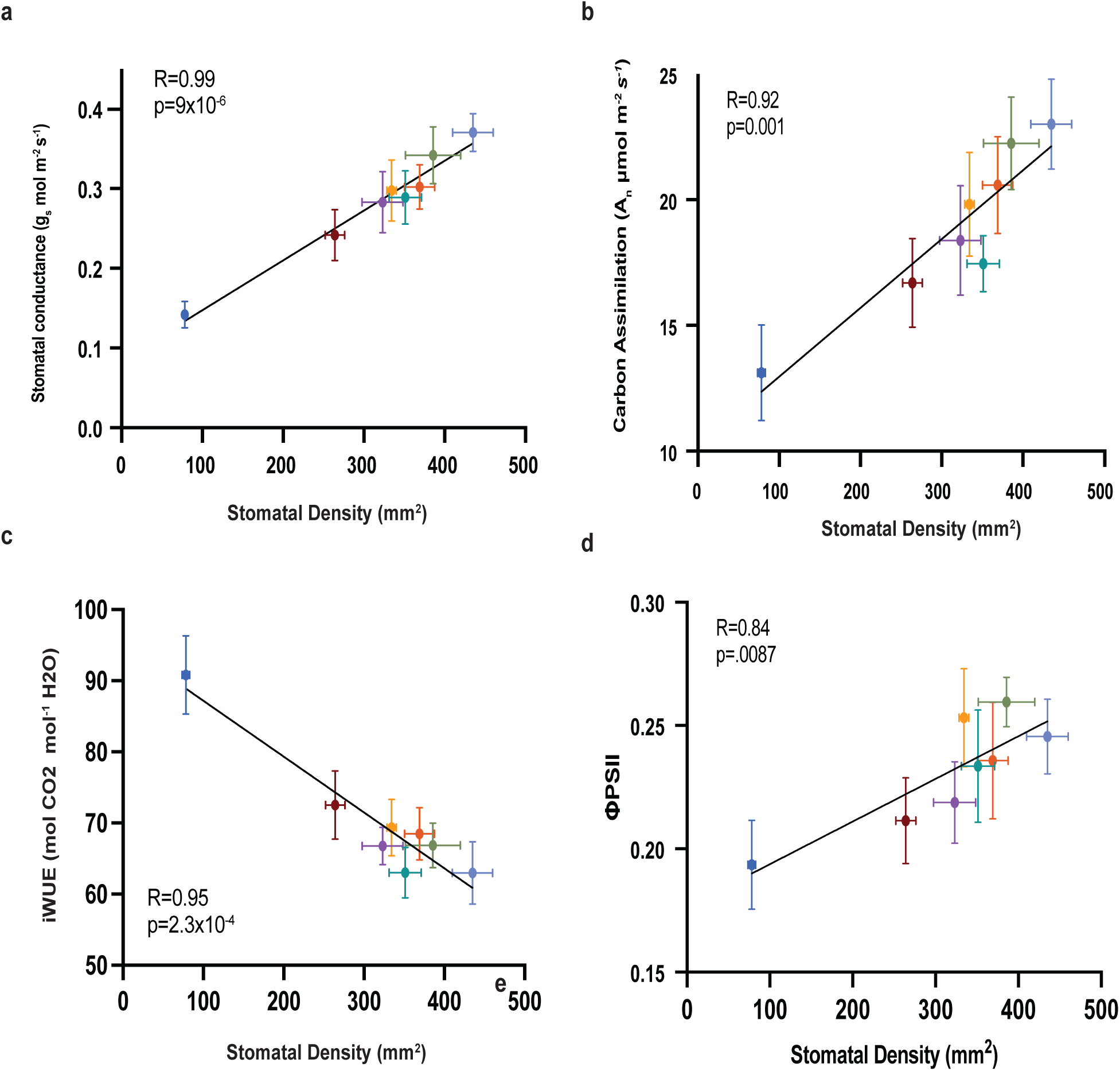
Stomatal morphological variation corresponds to gas exchange variation in a near-isogenic panel. Linear regression of stomatal density and (a) stomatal conductance, (b) carbon assimilation (c) intrinsic water-use efficiency, and (d) ΦSII. Correlation coefficient (R) and p-value of each correlation are noted in each panel. Mean and standard error of the mean are reported.

### Fluctuating light assay reveals dynamic gas exchange phenotypes across the stomatal density panel

To further investigate the relationship of stomatal variants to physiology in a context that more closely resembles field conditions, gas-exchange phenotypes were measured under a dynamic light regime (Figure 3a). Similar to the steady-state phenotypes reported in Figure 2, stomatal conductance and carbon assimilation maintain a positive association with stomatal density and a negative association with iWUE (Figure 3b, 3c). However, the cumulative iWUE, calculated from the area under the curve in dynamic light conditions, was lower than the steady state measurement in all genotypes. Additionally, the difference among genotypes in dynamic light regime was less pronounced (Figure 3d). The greatest discrepancy between measurements of cumulative iWUE according to assay type was recorded in *stomagen* where stomatal density is lowest and stomatal size is greatest (Figure 3d). To further resolve this difference in steady-state and dynamic light conditions, the rate of stomatal conductance reduction in response to transition from high light to low light was measured (Figure 3a). A one-phase decay model was fit to the stomatal conductance data immediately following the low light transition for each biological replicate, and rate constants were derived as a proxy for stomatal closure rate. Low density, large stomata lines *stomagen* exhibited the slowest closure rate constant, with variation in closure rate constants generally varying with stomatal density among other genotypes (Figure 3e).

**Figure 3.**
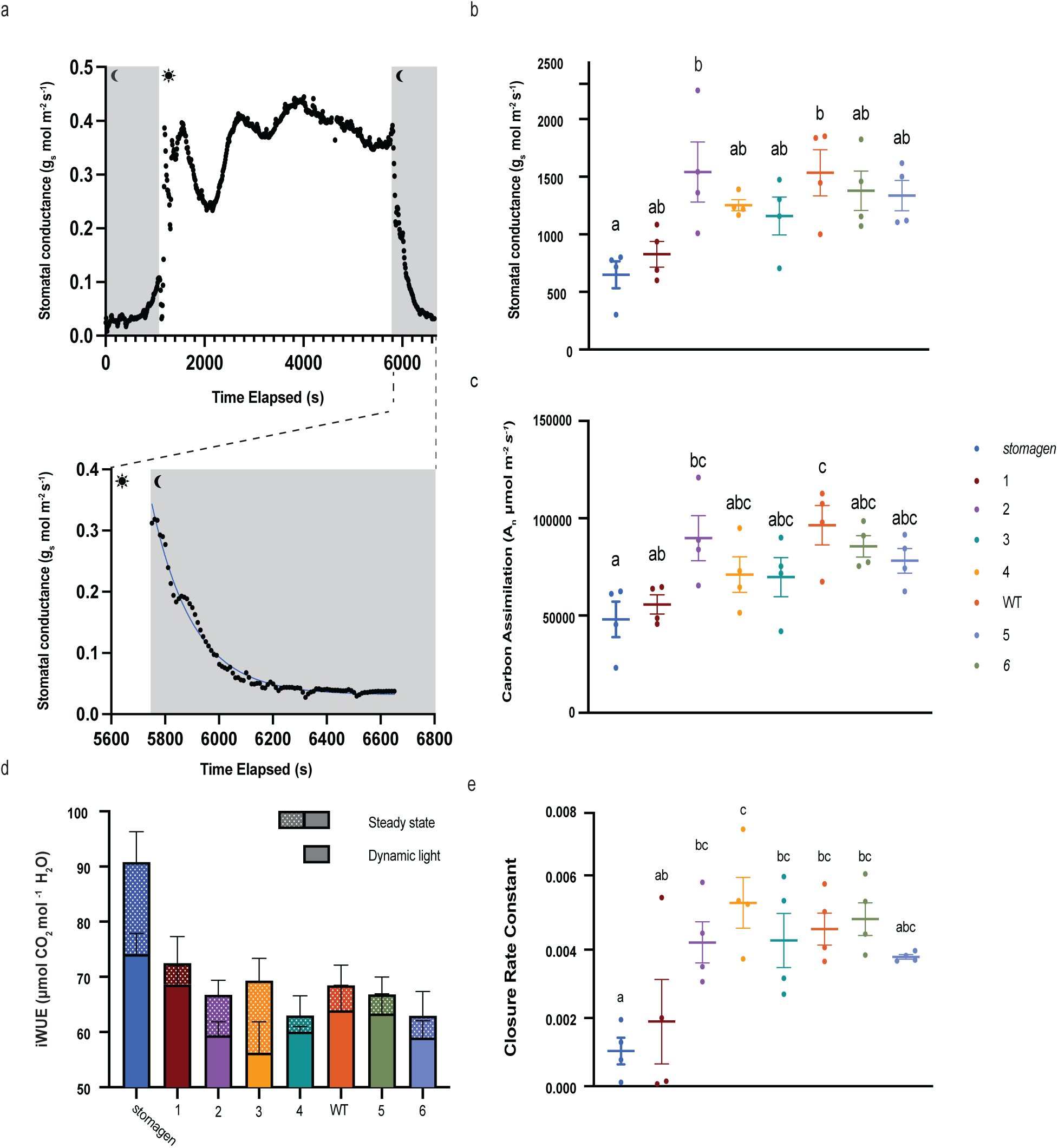
Physiological response to dynamic environmental conditions. (a) Representative stomatal conductance response curve to fluctuating light with gray blocks representing low light and white blocks representing high light. The lower inset is a representative one-phase decay curve fit to the second low light region of the curve. Dotplot of (b) cumulative stomatal conductance and (c) cumulative carbon assimilation across the entire regime, as calculated by area under the curve. (d) A stacked barplot of iWUE calculated from steady-state and dynamic light conditions. Solid blocks refer to dynamic light conditions and patterned blocks in addition to solid blocks reflect steady-state values. (e) Closure rate constant among alleles assayed in fluctuating light regimes, derived from the one phase decay curve fit to the stomatal conductance data following return from high light to low light. In the dotplots, each dot represents a biological replicate with bars indicating mean and standard error of the mean. Letters indicate a significant difference between means (P<0.05, one-way ANOVA Tukey HSD post-hoc test). In the barplot, the mean is represented with error bars showing standard error of the mean.

### Capacity for physiological and developmental response to abiotic stress differs among promoter alleles

The response of stomatal conductance in stomatal morphological variants under vegetative drought was also assayed. The variation identified in well-watered conditions was nearly eliminated after imposing a 5-day vegetative drought (Figure 4a,b,c). All alleles besides *stomagen* exhibited an overall reduction in stomatal conductance in response to vegetative drought (Figure 4c).

**Figure 4.**
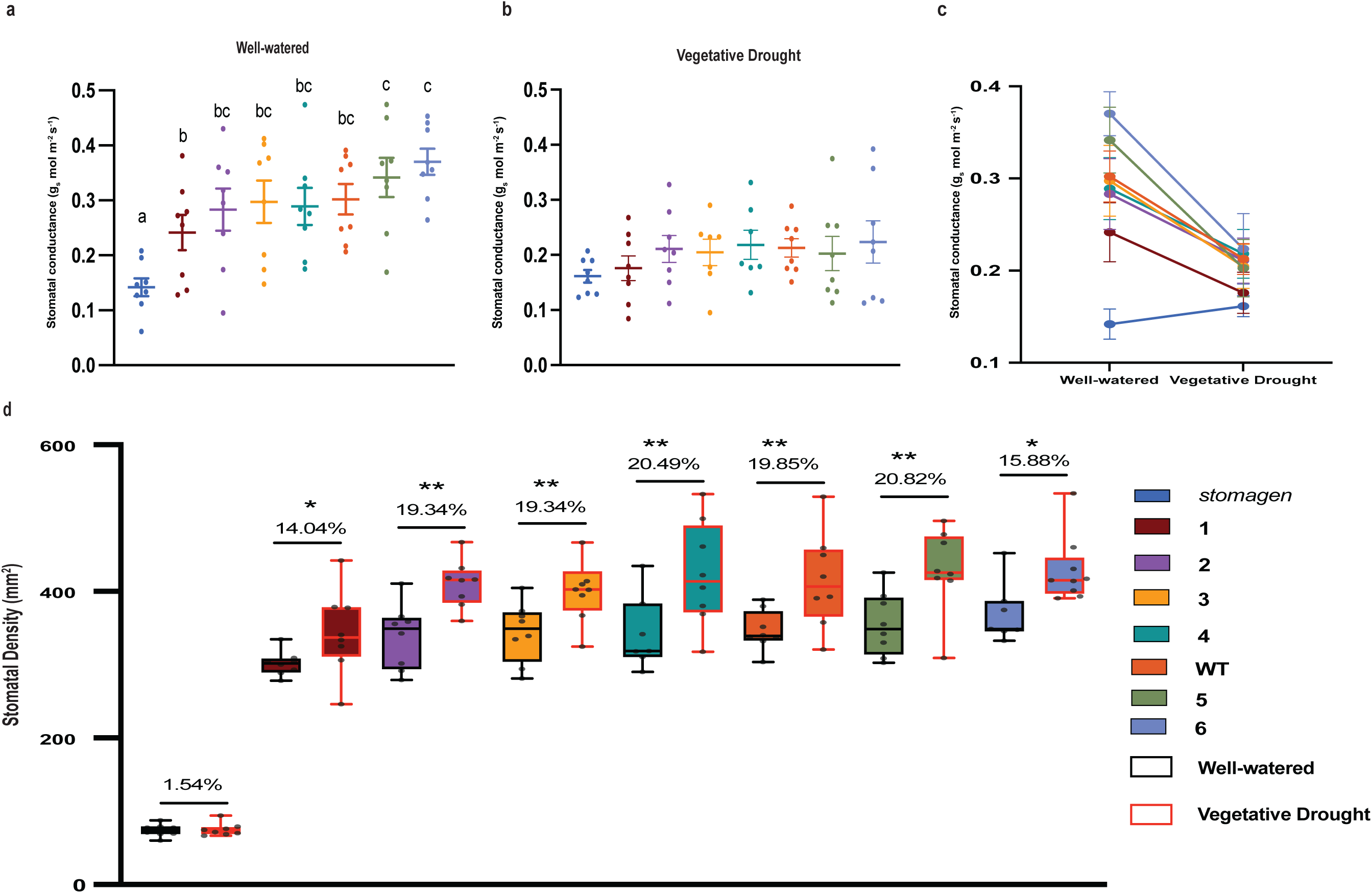
Physiological and developmental response of stomatal variants to vegetative drought. Dotplot of stomatal conductance in (a) well-watered plants and (b) after a 5-day vegetative drought. (c) A reaction normalization plot of stomatal conductance in each watering regime. (d) Changes in stomatal density in response to vegetative drought. The percent increase of vegetative drought stomatal density relative to well-watered is reported above each genotype. In the dotplots, each dot represents a biological replicate with bars indicating mean and standard error of the mean. Letters indicate a significant difference between means (P<0.05, one-way ANOVA Tukey HSD post-hoc test). In the box-and-whisker plot, the center horizontal indicates the median, upper and lower edges of the box are the upper and lower quartiles and whiskers extend to the maximum and minimum values within 1.5 interquartile ranges. Barplot shows means and error bars represent standard error of the mean. In (f) black and red outlines of represents well-watered and vegetative drought, respectively. * represents a p-value <0.1, <0.05, and ** represents a p-value <0.05 (Student’s t-test).

Whereas *stomagen* maintained a near consistent overall stomatal conductance regardless of drought status, other stomatal morphological variants exhibited significant declines in stomatal conductance (Figure 4b,c). A similar drought assay was imposed on a separate cohort of plants to monitor drought-responsive changes in stomatal density. A moderate vegetative drought resulted in stomatal density increases in the wild type background (Figure 4d). This finding is consistent with previous studies where a vegetative drought was applied to a grass species^37,38^. The *stomagen* mutant, however, exhibited very limited stomatal density reprogramming response after drought (Figure 4d).

Most *OsSTOMAGEN* promoter alleles exhibited a wild-type increase in stomatal density of 20% after abiotic stress (Figure 4d). However, in alleles 1 and 6, the capacity for drought-responsive stomatal density reprogramming was slightly diminished (Figure 4d).

*OsSTOMAGEN* transcript abundance after abiotic stress treatment was analyzed to discern the effect of promoter edits on expression. A drought-responsive transcriptional increase of *OsSTOMAGEN* in wild-type developing leaves was consistent with the observed increase in stomatal density (Extended Data 3). Likewise, promoter allele 1 also exhibited an increase in *OsSTOMAGEN* transcript abundance, despite exhibiting attenuated stomatal density reprogramming. No *OsSTOMAGEN* expression increases after vegetative drought were detected in any other promoter allele. Consistent with previous *OsSTOMAGEN* expression data, we observed limited co-linearity between expression levels and phenotype in response to vegetative drought, despite a consistent stomatal density increase among most promoter alleles (Extended Data 1d).

## Discussion

Cis-regulatory editing approaches provide the opportunity to rapidly generate novel quantitative variation. Application of this approach to *OsSTOMAGEN*, a positive regulator of stomatal density, produced an array of promoter alleles representing 70% to 120% of wild-type stomatal density. A multiplexed guide RNA design approach leveraging publicly available bioinformatic datasets enabled the selection of guides that ultimately had a substantial impact on phenotypic outcomes. Notably, a higher stomatal density was observed in two edited promoter alleles.

Hypermorphic phenotypes generated by targeted and unbiased promoter editing have been associated with large chromosomal rearrangements^39,40^. Here we report two hypermorphic alleles produced by proximal indels alone, suggesting that the capacity to generate gain-of-function variants may be possible without larger perturbation and may be highly target-gene dependent.

One of the unique attributes of these two alleles is a fully unedited region that corresponds to a CNS peak shared with the C3 grasses *B. distachyon* and *H. vulgare.* The transition of C3 to C4 photosynthesis includes a marked reduction of stomatal density linked to reductions in *STOMAGEN* expression^9^. It is noteworthy that the only two alleles that maintain the entire conserved region among C3 Poaceae family members, with edits in other promoter regions, result in a hypermorphic phenotype. It is possible that this C3-specific CNS peak contributes positively to the stomatal density phenotype, whereas CNS peaks conserved within the whole family regulate basal levels of stomatal development. Cis-regulatory editing programs may benefit from editing divergent cis-regulatory elements towards an increased understanding of how novel *cis*-regulatory elements contribute to novel trait emergence.

To understand the transcriptional responses of edited promoter alleles, the transcript abundance of *OsSTOMAGEN* was measured across various developmental stages and abiotic stresses. There were some differences in *OsSTOMAGEN* expression levels among promoter alleles, however, these transcriptional differences did not correspond to correlative stomatal density outcomes. This finding is consistent with some previous applications of promoter editing where expression levels of targeted genes were not explanatory of phenotypic outcomes^16,41,42^. In examples where expression levels do not predict phenotypes, mis-regulation of expression of targeted genes may occur in highly discrete spatial or temporal zones that may diverge from native gene expression and can be difficult to capture. Thus, gene expression may be insufficient as a proxy for phenotype.

Beyond stomatal density, promoter alleles of *OsSTOMAGEN* had implications on stomatal size. There is a well-documented inverse relationship of stomatal density and stomatal size across many species including rice ^2,8,43–45^. This trend remained among the panel of promoter alleles (Supplemental Figure 4), supporting a previously hypothesized role of OsSTOMAGEN in mediating stomatal size in addition to density^46^. Deviation of gene-edited guard cell lengths from the expected inverse relation may thus be similarly affected and confounded by the observed stochasticity in *OsSTOMAGEN* expression across tissue types and stresses (Extended Data Figures 1,2,3).

Our stomatal density panel enhanced the resolution of density versus gas exchange phenotypes. A strong, positive, linear, relationship between stomatal density and stomatal conductance and carbon assimilation were established, consistent with previous reports. Likewise, a strong, negative linear association of stomatal density and iWUE was found ^2,8,21,47^. Interestingly stomatal density was found to have a positive, linear relationship with ΦPSII. Transpiration via stomata plays an essential role in evaporative cooling under peak diurnal temperatures and irradiances. Long-term or repeated exposure to heat stress, exacerbated by low stomatal densities, may contribute to this relationship of density and ΦPSII^48^.

A fluctuating light assay revealed additional structure-function relationships beyond traditional steady-state measurements. Steady-state measurements likely overstate the gains in water-use efficiency of reduced density lines, as shown by the decrease in iWUE differences under dynamic, fluctuating light. Discrepancies may result from lowered rates of stomatal closure in low density, larger stomata lines. These findings may reconcile previous data that show similar levels of water loss from lines with moderate and strong reductions in stomatal density. Engineering stomatal density reductions without effecting size and/or kinetics may thus be promising mechanism to generate WUE improvements in dynamic field conditions.

Our CRISPR near-isogenic lines (cNILs) provide a major advantage in overcoming previous limitations of studies relying on natural variation in stomatal density phenotype across diverse germplasm, resolving discrete structure-function relationships. Within this study, the cNIL approach resolved unexpected correlations between stomatal density and size, ΦPSII, and fluctuating light gas exchange response—all traits that can be influenced by genetic heterogeneity. Quantitative gene editing can thus expand our basic understanding of other polygenic traits as well, such as the relationships of root lengths to drought tolerance, plant height to yield, or trichome density to pathogen resistance, among many other possibilities^42^.

The cNIL panel also enhanced our ability to interrogate developmental plasticity under drought stress, which is known to cause stomatal density reprogramming^37,38,49^. Consistent with a previous report of stomatal density shifts in drought stress, we find moderate increases in wild-type stomatal density in the drought regime. Stomatal closure induced by drought stress may result in less evapotranspirative cooling, leading to warmer leaf temperatures and overall greater heat stress ^2,8^. Stomatal density increases may be a heat stress response to increase rates of cooling^37,38^. Stomatal density increases correlated with increases in *OsSTOMAGEN* expression in the wild type. Contrary to expectations, mis-regulation of *OsSTOMAGEN* in most gene-edited alleles under vegetative drought conditions suggest additional nuance in environmentally responsive gene expression and phenotype. In this case, stomatal density reprogramming capacity during abiotic stress was lower in the lowest (Allele 1) and greatest (Allele 6) stomatal density alleles. There are however no apparently unique edits common to these two alleles. The high extent of misexpression may be linked to a reduced capacity for environmentally-responsive transcriptional reprogramming of *OsSTOMAGEN* Aside from differences in development, we also describe greater magnitude changes in drought-responsive stomatal conductance as indicated by the reaction normalization plot (Figure 4c). Varied physiological and developmental responses to vegetative drought among promoter alleles offers greater insights to allele compatibility in environments with varied water availabilities.

Overall, we demonstrate the diverse, complex implications of promoter editing on trait variation in dynamic light and water environments. Applications of promoter editing for improved crop improvement must consider pleiotropy of edits in dynamic environments inherent to nearly all production landscapes. There is also an opportunity to expand promoter editing applications for improving environmental response outcomes. For example, promoter editing has been leveraged as a tool to prevent upregulation of susceptibility loci during pathogen invasion^50^. Tuning environmental response by gene editing of cis-regulatory elements may also be a desirable approach for other traits achievable by promoter editing.

The developmental and physiological insights generated by cis-regulatory element editing can be further leveraged into crop improvement. Rice production spans highly variable environmental conditions, especially with regards to water availability. The morphological diversity generated and the corresponding physiology characterized can be matched with the most suitable environmental (Figure 5). For instance, low density lines with greater iWUE may be most appropriate in field sites with low water availability. Whereas, in water replete conditions, higher density lines with greater rates of carbon assimilation may be best suited. In all rain-fed production schemes water availability is variable. In these conditions, selecting alleles that exhibit full wild-type capacity for stomatal density reprogramming is likely desirable for maintaining fitness. Elite cultivars confined to narrow geographies due to limitations in a discrete set of traits could theoretically be optimized by promoter editing for compatibility to a much broader set of environments. Taken together, our data offer, an improved understanding of the opportunities of cis-regulatory element editing in generating quantitative trait variation for broad and dynamic environments, and an expansion of promoter editing as a tool for: generating hypermorphic trait variation without large chromosomal rearrangements, establishing near isogenic panels for assessments of discrete structure-function relationships, and generating expansive trait diversity compatible with similarly expansive environments.

**Figure 5.**
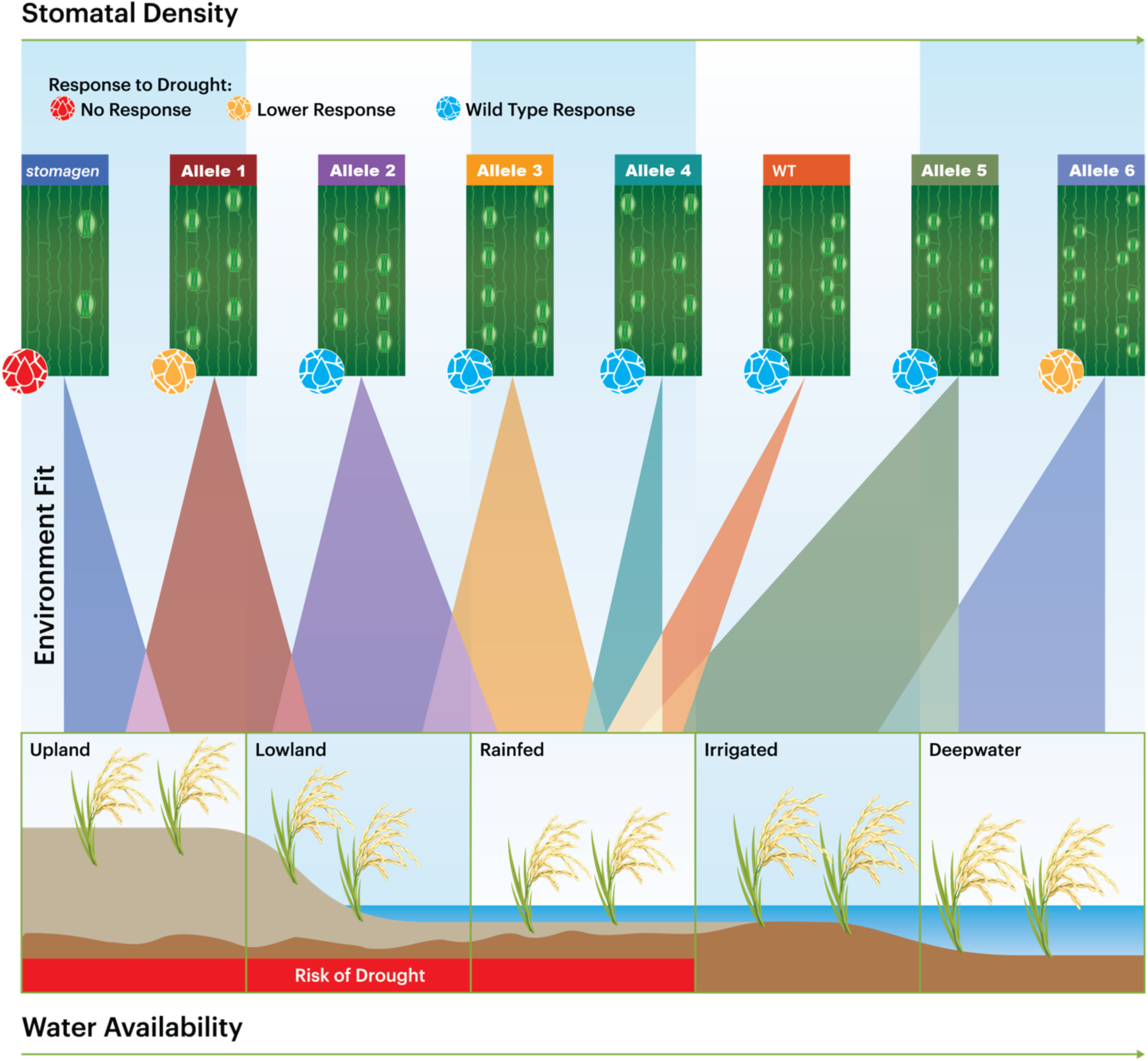
Stomatal morphological variation can be matched to broad and dynamic production environments. Rice is produced in diverse environments with varying water availabilities and fluctuations. Variation generated by gene editing can be matched to variation in environments. Each stomatal variant is represented by a graphic of stomatal density and size. Colored spotlights indicate potential environmental range of each allele according to stomatal characteristics including stomatal density, stomatal conductance, and responsiveness to abiotic stress represented by red, yellow, and blue water droplet icons.

## Acknowledgements

We thank Christina Winstrom and all the greenhouse staff who helped care for our plants. We also acknowledge Bastian Minkenberg for cloning the guide construct and Glenn Ramit for support in producing figures. DP-T was supported by the Berkeley Fellowship and the NSF Graduate Research Fellowship Program (Grant DGE 1752814). KKN is an investigator of the Howard Hughes Medical Institute.

## Author Contributions

NGK developed project concept and coordinated research efforts. MT transformed and regenerated rice plants in the lab of MJC with technical support from MJC. Rice stomatal density phenotyping was led by NGK with support from AGC, LL, and SAL. qPCR experiments were completed by NGK, AGC, LL, and SAL. Physiological experiments were completed by NGK with technical advising from DPT. LICOR equipment was made available by KKN. All work was completed in the lab of BJS. The manuscript was drafted by NGK and edited by DPT.

## Notes

### Competing Interest Statement

The authors have declared no competing interest.

