## Supplemental Figures and Tables for "Editing cis-regulatory elements towards generating rice stomatal morphological variation for adaptation to broad and dynamic environments"

Extended Data 1:

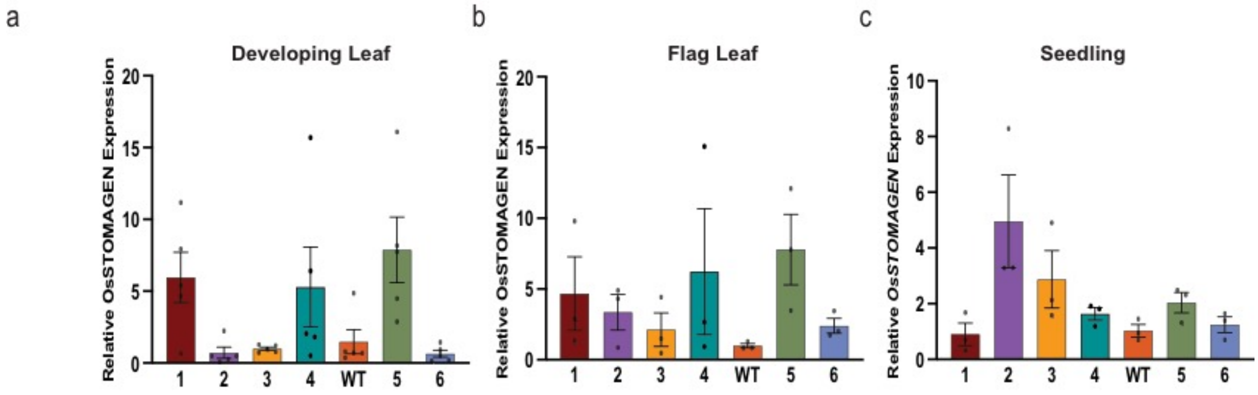


Barplot of the absolute normalized relative expression of *OsSTOMAGEN* of each allele in (a) developing leaves, (b) flag leaves and (c) seedlings normalized to WT expression in each tissue type. In the barplots mean is represented with error bars showing standard error of the mean.

Extended Data 2:

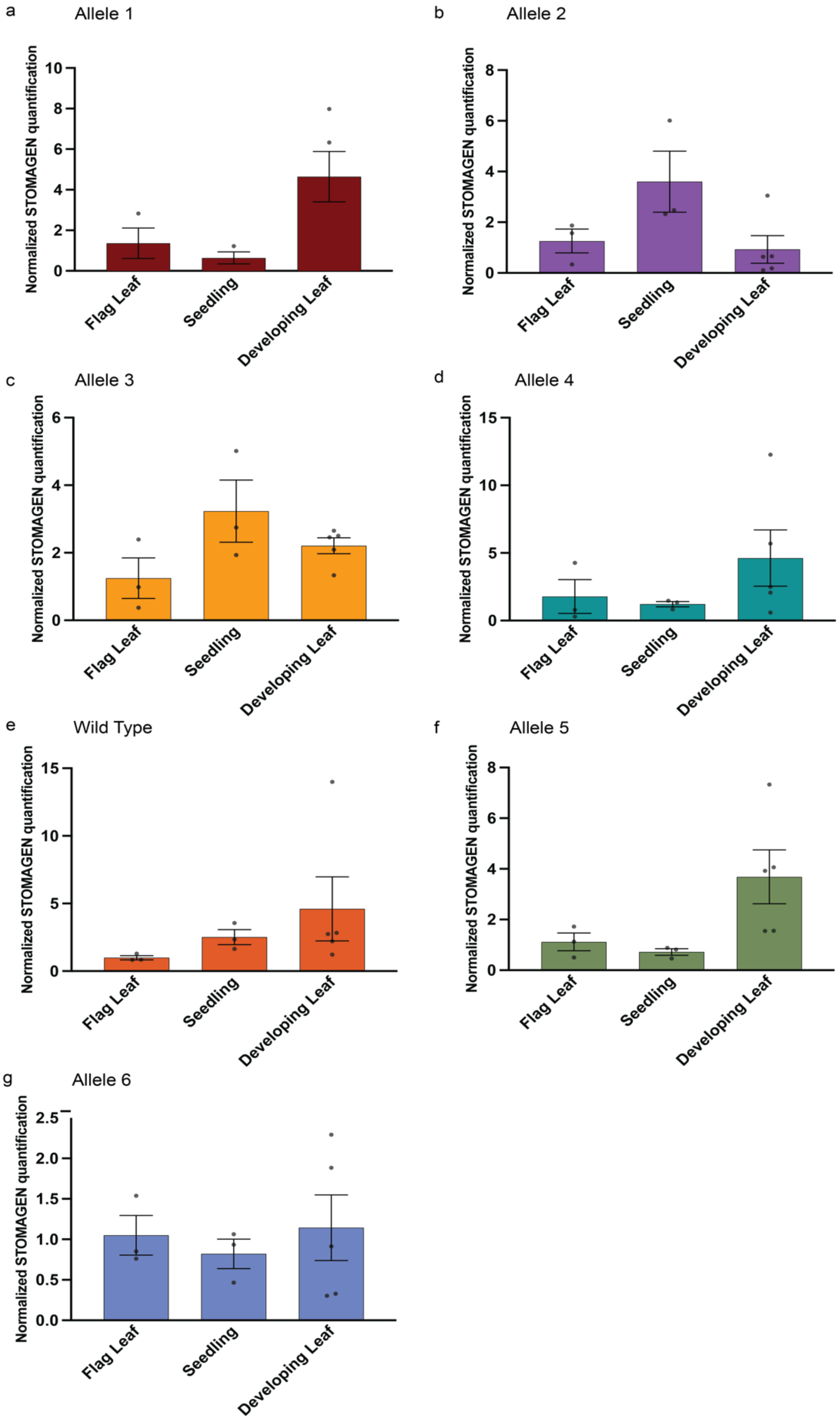

Extended Data 2| Relative Expression of *OsSTOMAGEN* in varying tissues within each promoter allele

A comparison of *OsSTOMAGEN* transcript abundance among flag leaves, seedlings, and developing leaves in (a) Allele 1 (b) Allele 2 (c) Allele 3 (d) Allele 4 (e) Wild type (f) Allele 5 (g) Allele 6. In each genotype values are relative to flag leaf wild-type expression and normalized to the average of two housekeeping genes. Barplot shows means and error bars represent standard error of the mean.

Extended Data 3:

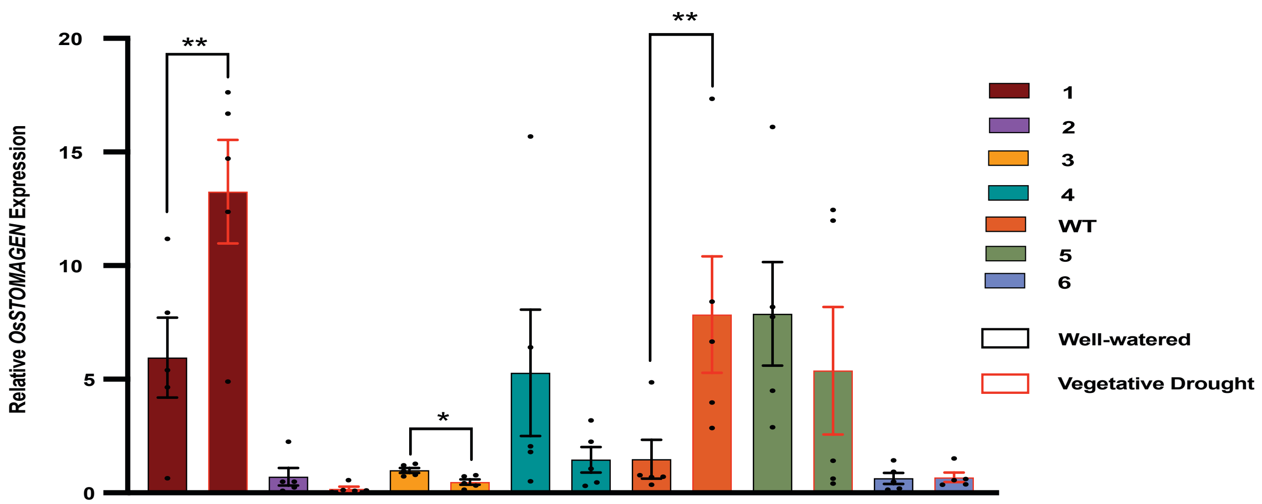


Extended Data 3|

*OsSTOMAGEN* expression in each allele in well-watered and vegetative drought. Barplots mean is represented with error bars showing standard error of the mean. Asterisks represent a significant difference in expression relative to wild type (P<0.05, one-way ANOVA Tukey HSD post-hoc test). Black and red outlines of represents well-watered and vegetative drought, respectively. * represents a p-value <0.1, >0.05, and ** represents a p-value <0.05 (Student's t-test).

Supplemental Figure 1:

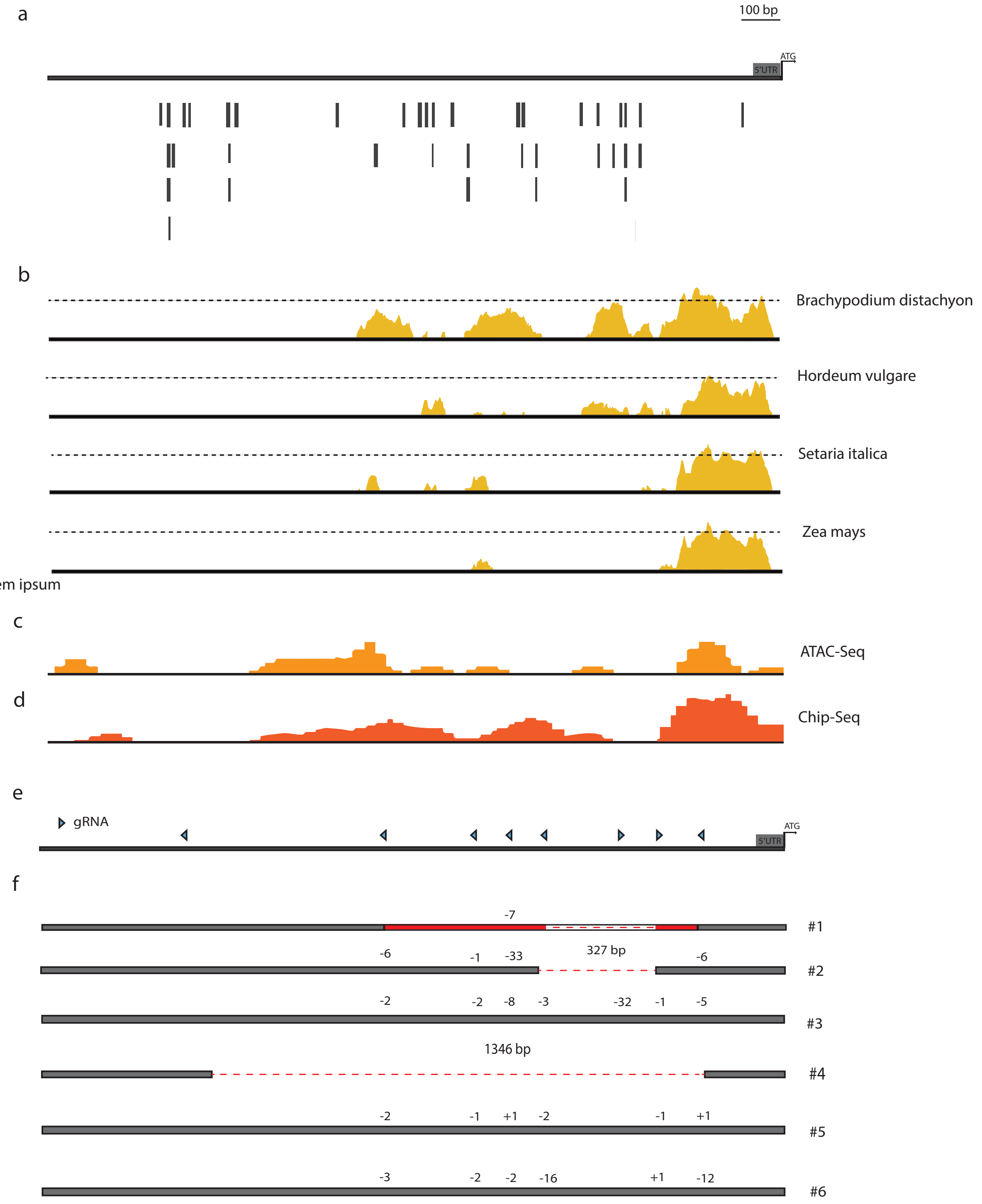

Supplemental Figure 1| Rational guide design approach for targeting the promoter of *OsSTOMAGEN*

(a) Putative transcription factor binding sites using a 99% identity threshold annotated with reference to translation start site (b) mVISTA plot displaying conserved non-coding sequences among evolutionarily dispersed Poaceae family members. The dashed line represents a 75% conserved threshold. Peaks displayed represent regions of minimum 50% conservation. Poaceae family members are arranged from most similar to most distant relatives (c) ATAC-seq data extracted from RiceENCODE is shown for the promoter region of *OsSTOMAGEN* (d) ChIP-seq data for H3K27ac, extracted from RiceENCODE database. (e) A summary of the positions and orientations of the guide sequences used to target the promoter of *OsSTOMAGEN.* Each blue triangle represents an individual guide, with the triangle pointing towards the 3' NGG site. (f) An overview of each unique allele generated. Large deletions are represented by red dashed lines and indels at each guide site are denoted by (+) to indicate insertions or (-) for deletions alongside a number representing total quantity of base pairs associated with each indel. Red blocks indicate inverted sequences. Each allele is labeled with a unique number.

Supplemental Figure 2:

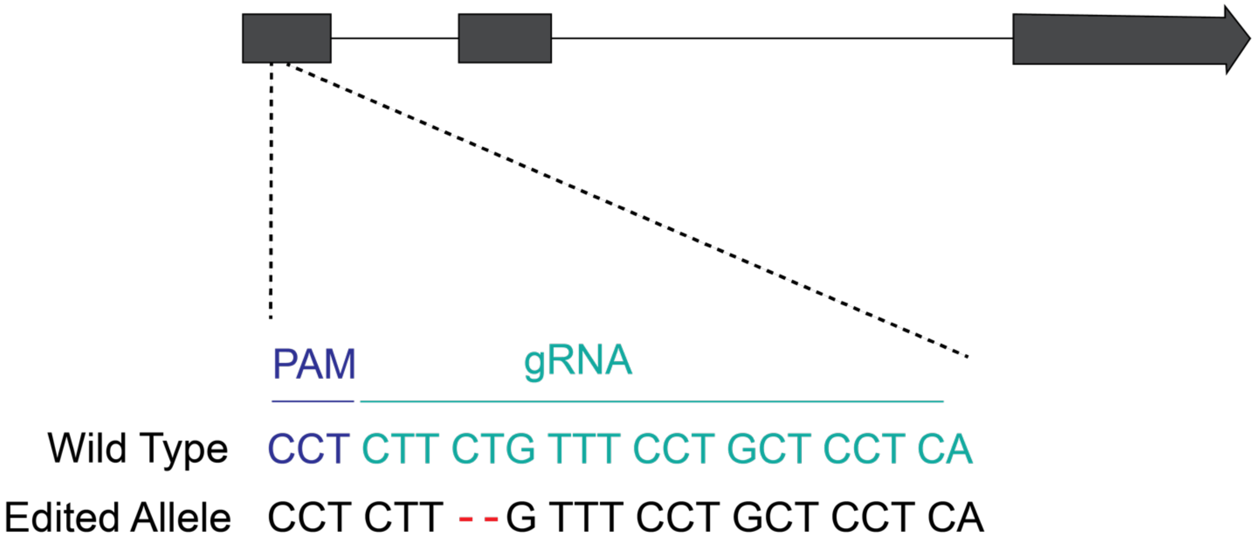


Supplemental Figure 2| Genotype of edited *stomagen* allele

The gene model of *OsSTOMAGEN* with the location of the CRISPR/Cas9 guide RNA indicated in blue. The unique edits generated by CRISPR/Cas9 are shown in red.

Supplemental Figure 3:

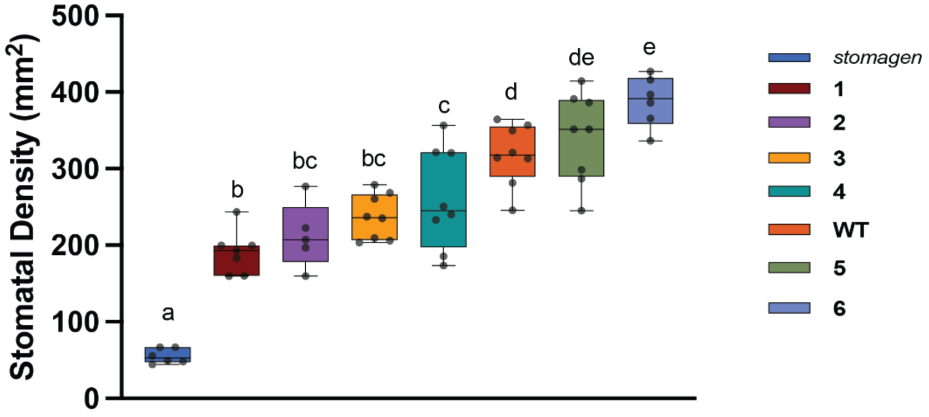


Supplemental Figure 3 | Stomatal density variation of promoter alleles grown in growth chamber

a) Box-and-whisker plot of the stomatal density of each allele assayed. In the box-and-whisker plot, the center horizontal indicates the median, upper and lower edges of the box are the upper and lower quartiles and whiskers extend to the maximum and minimum values within 1.5 interquartile ranges. Each dot represents a biological replicate. Letters indicate a significant difference between means (P<0.05, one-way ANOVA Tukey HSD post-hoc test). Plans were grown in chambers at 28 °C for day-length periods of 16 h in 400 $\mu$mol photons/$m^2$/s of light and 80% relative humidity.

Supplemental Figure 4:

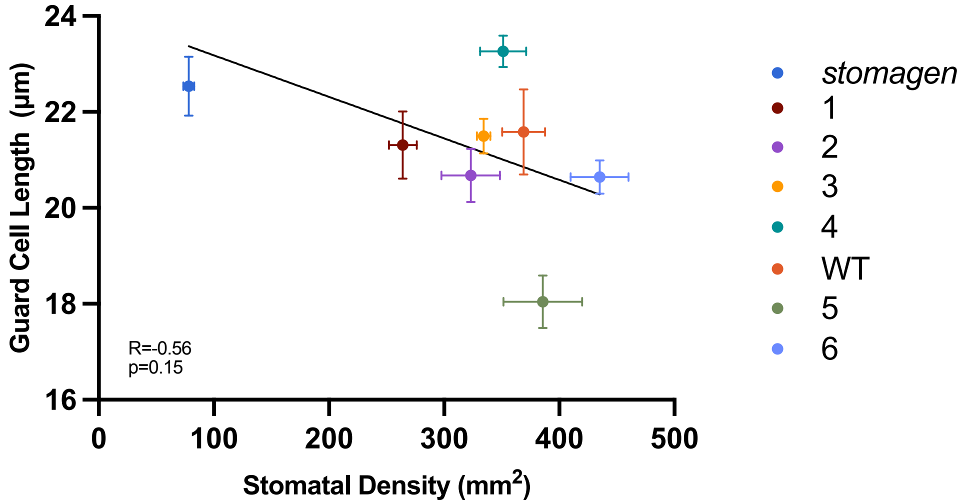


Supplemental Figure 4| Linear regression of stomatal density and guard cell length

Linear regression of stomatal density and guard cell length. The correlation coefficient (R) and p-value of the correlation is noted. Mean and standard error of the mean are reported.

Supplemental Figure 5:

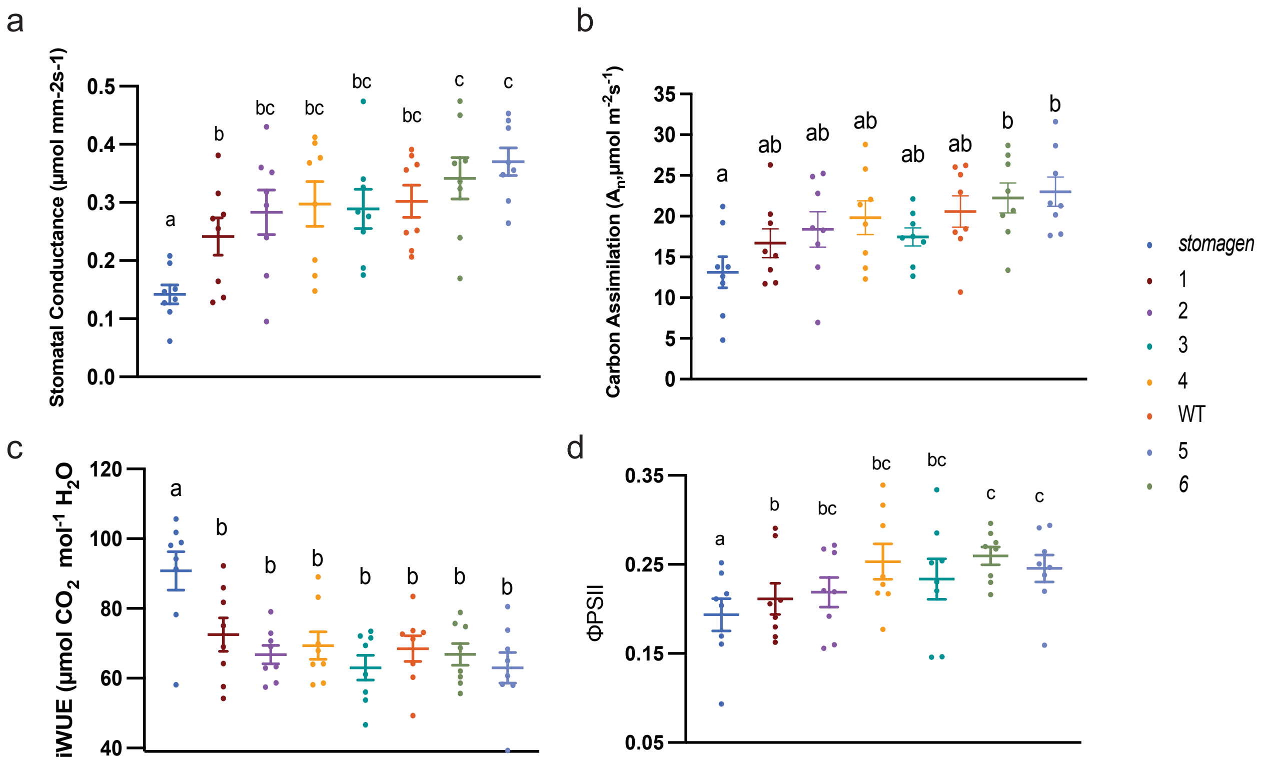


Supplemental Figure 5 | Well-watered greenhouse gas exchange measurements

Dotplot of (a) stomatal conductance (b) carbon assimilation (c) iWUE and (d) ΦPSII measured on each allele. Each dot represents a biological replicate with bars indicating mean and standard error of the mean. Letters indicate a significant difference between means (P<0.05, one-way ANOVA Tukey HSD post-hoc test

Supplemental Figure 6

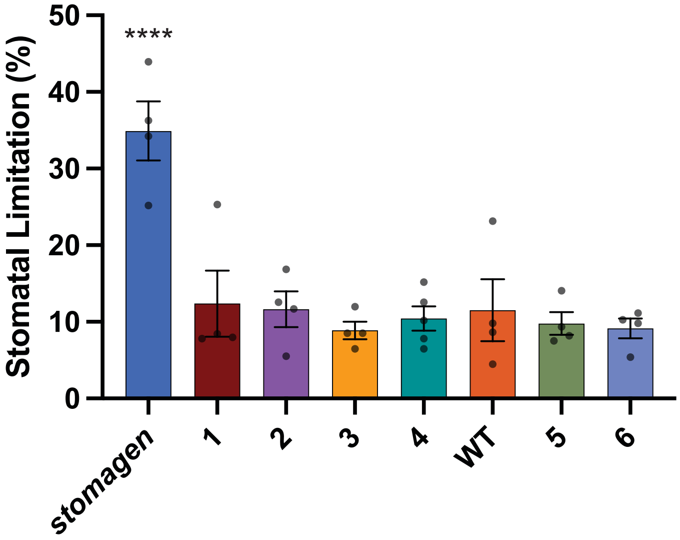


Supplemental Figure 6 | Stomatal limitation derived from A/Ci curves

A barplot of stomatal limitation percentages derived from A/$C_i$ curves generated on biological replicates of each allele showing mean and standard error of the mean. . Barplot shows means and error bars represent standard error of the mean.  **** represents a p-value <0.001

Supplemental Figure 7

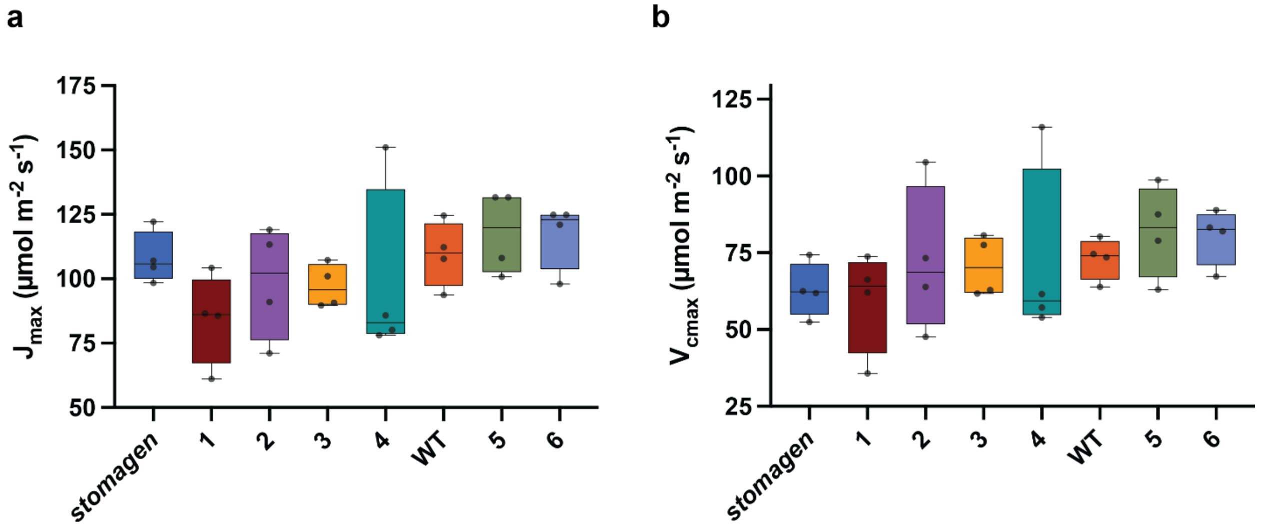


Supplemental Figure 7 | $J_{max}$ and $V_{cmax}$ of *OsSTOMAGEN* alleles

Box-and-whisker plot of (a) $J_{max}$ and (b) $V_{cmax}$ derived from A-Ci curves using Plantecophys package in R studio. In the box-and-whisker plot, the center horizontal indicates the median, upper and lower edges of the box are the upper and lower quartiles and whiskers extend to the maximum and minimum values within 1.5 interquartile ranges. Each dot represents a biological replicate Each allele is represented by four biological replicates.

Supplemental Figure 8

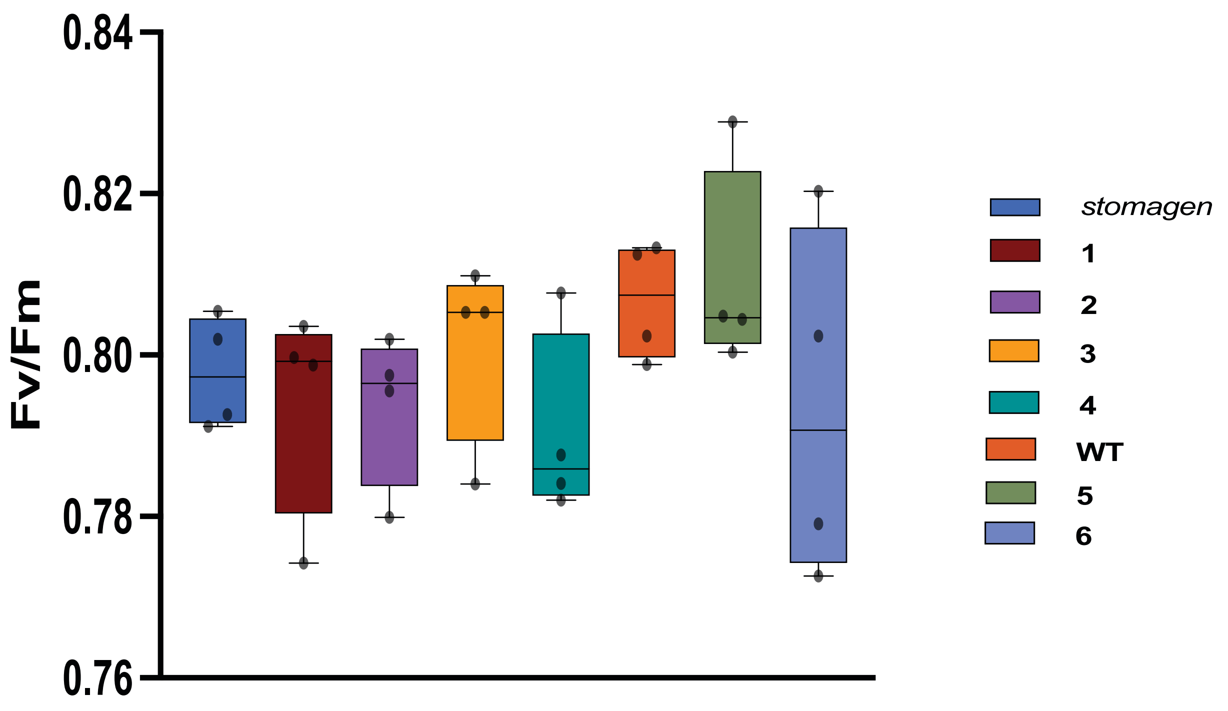


Supplemental Figure 8 | Chlorophyll fluorescence among promtoer alleles

Box and whisker plot of the Fv/Fm measurements of each promoter allele. In the box-and-whisker plot, the center horizontal indicates the median, upper and lower edges of the box are the upper and lower quartiles and whiskers extend to the maximum and minimum values within 1.5 interquartile ranges. Each dot represents a biological replicate.

Supplemental Figure 9:

**a**

*stomagen*

Replicate 1:

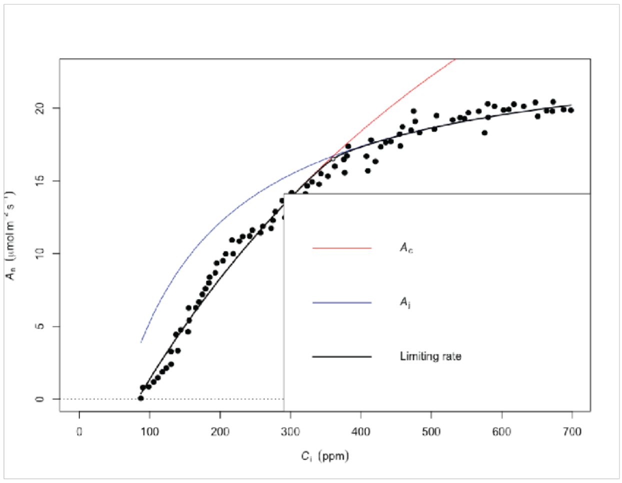


Replicate 2:

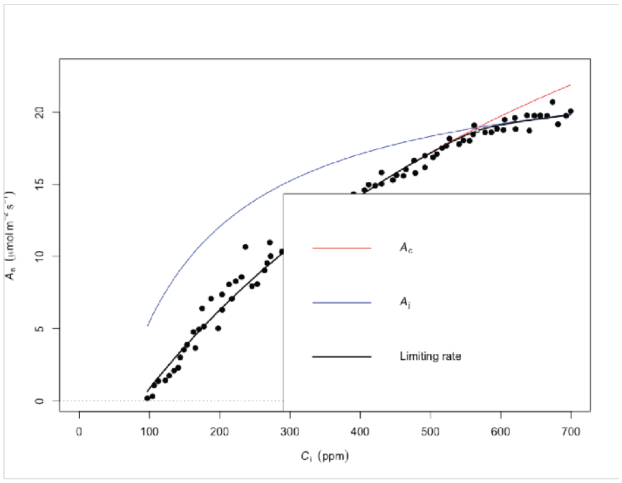


Replicate 3:

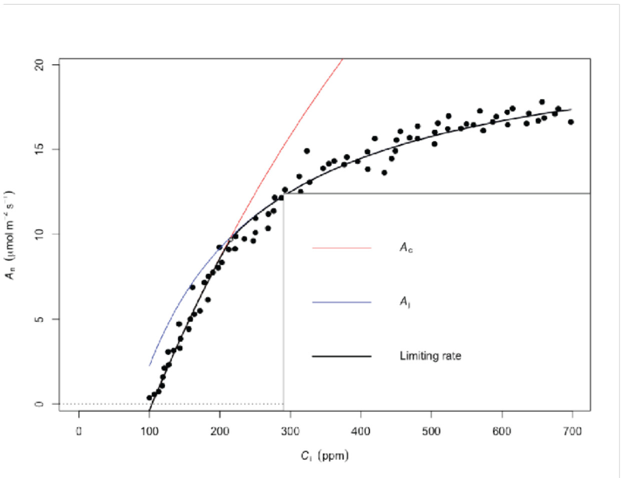


Replicate 4:

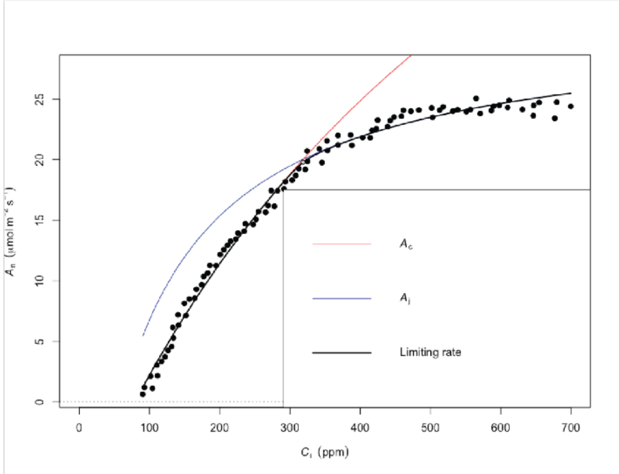

**b**

#### Allele 1

Replicate 1:

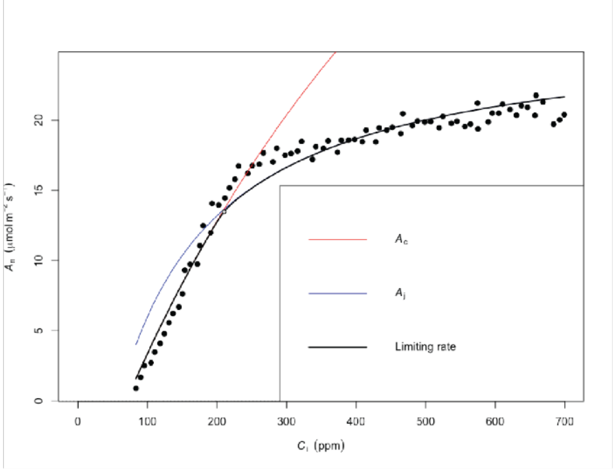


Replicate 2:

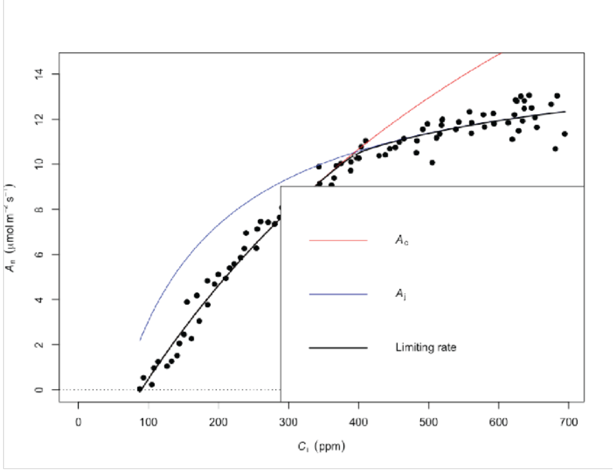


Replicate 3:

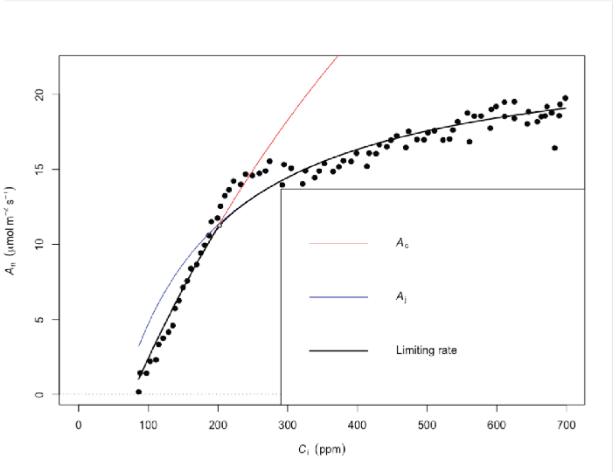


Replicate 4:

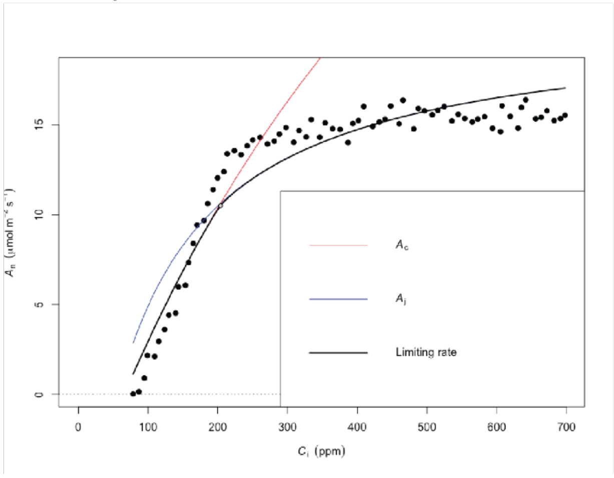

**c**

#### Allele 2

Replicate 1:

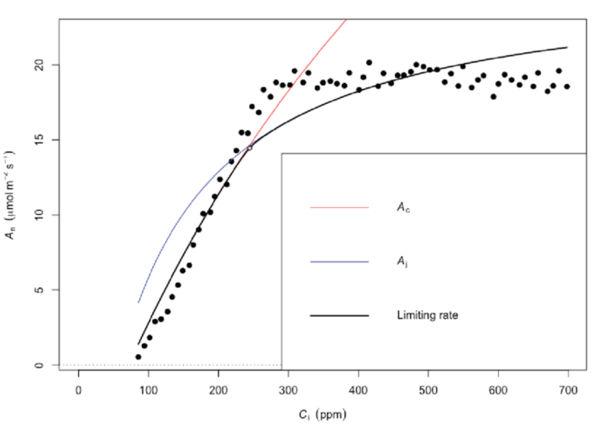


Replicate 2:

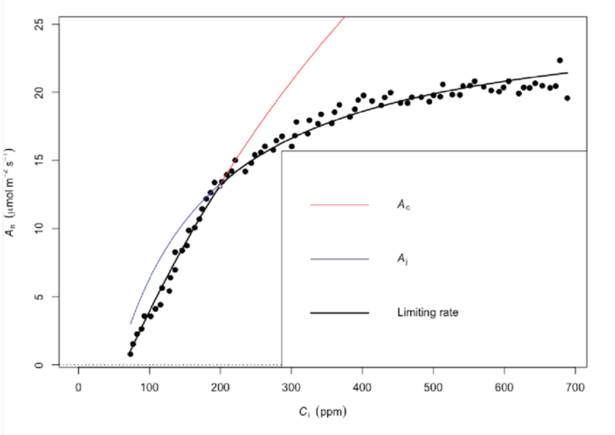


Replicate 3:

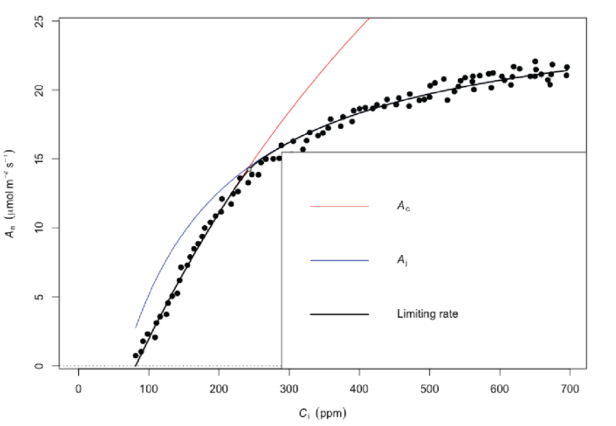


Replicate 4:

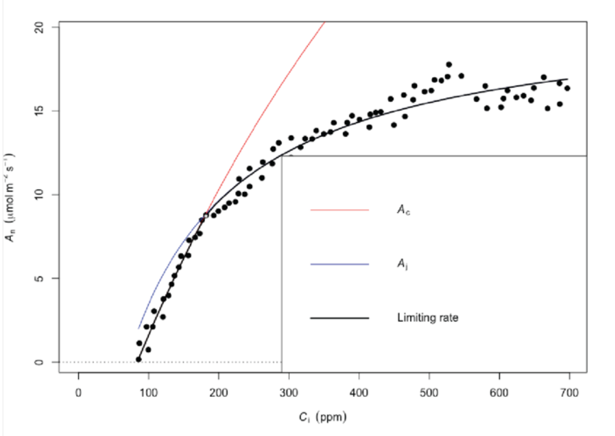

**d**

#### Allele 3

##### Replicate 1:

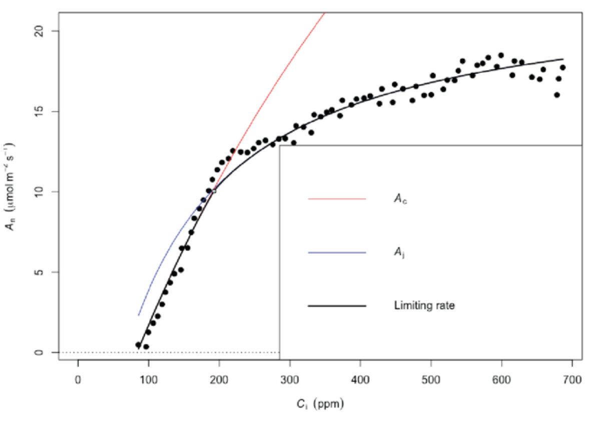


##### Replicate 2:

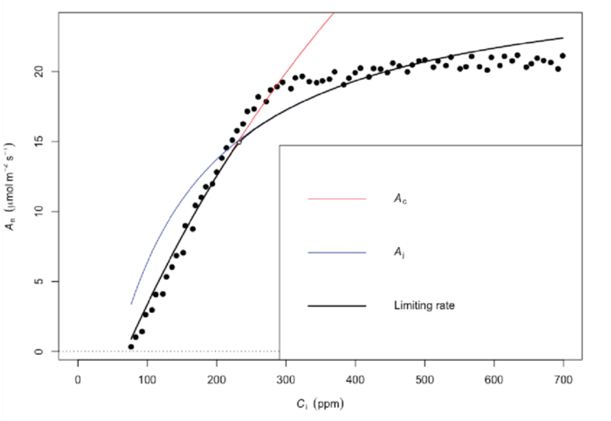


##### Replicate 3:

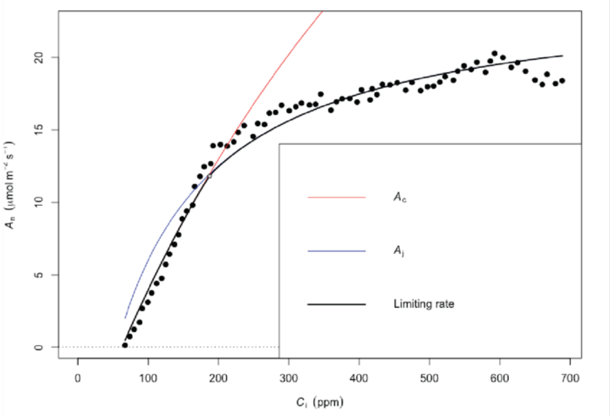


##### Replicate 4:

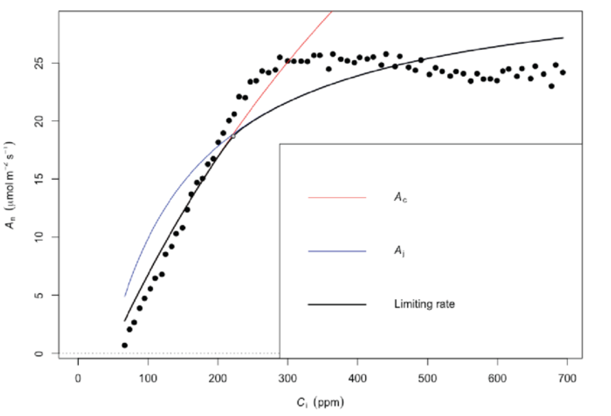

**e**

#### Allele 4

Replicate 1:

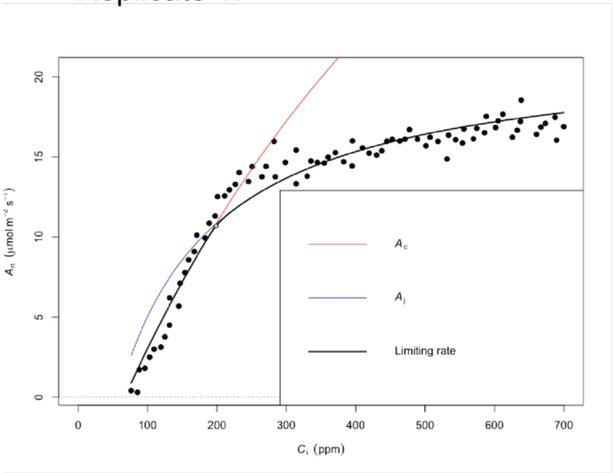


Replicate 2:

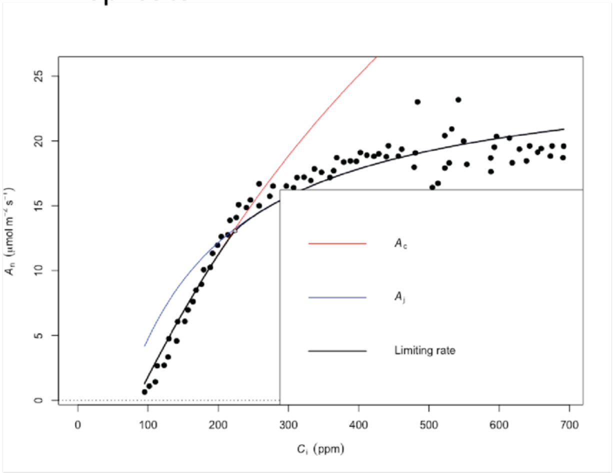


Replicate 3:

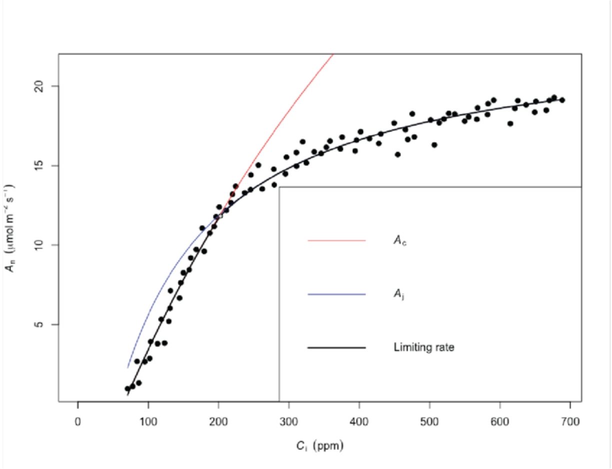


Replicate 4:

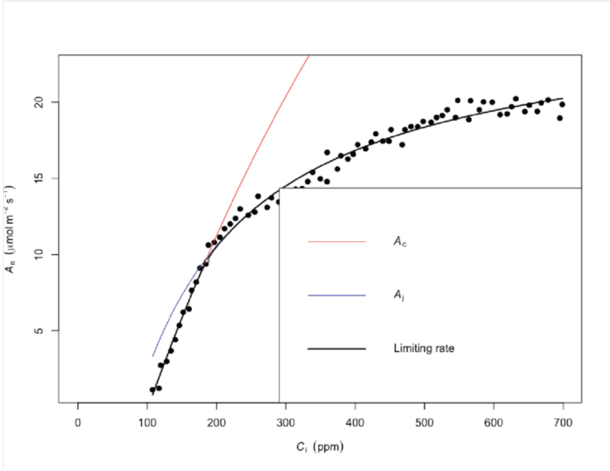

**f**

#### Wild type

Replicate 1:

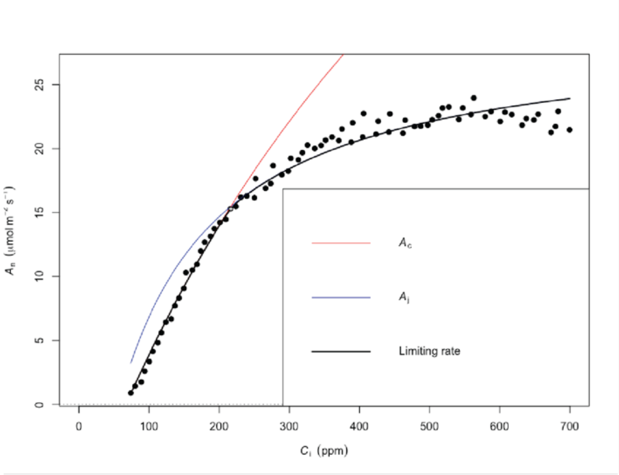


Replicate 2:

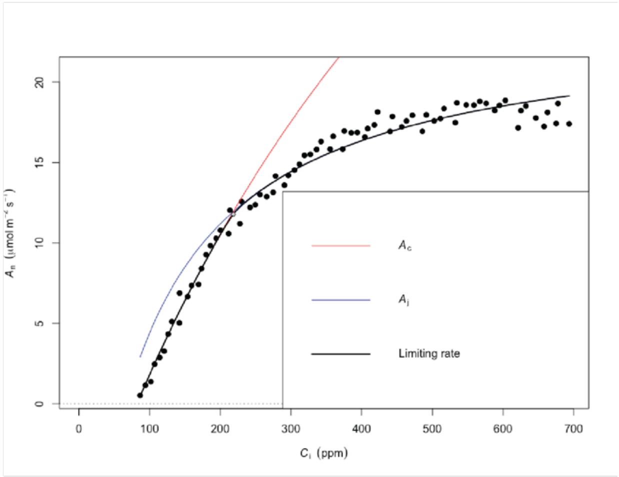


Replicate 3:

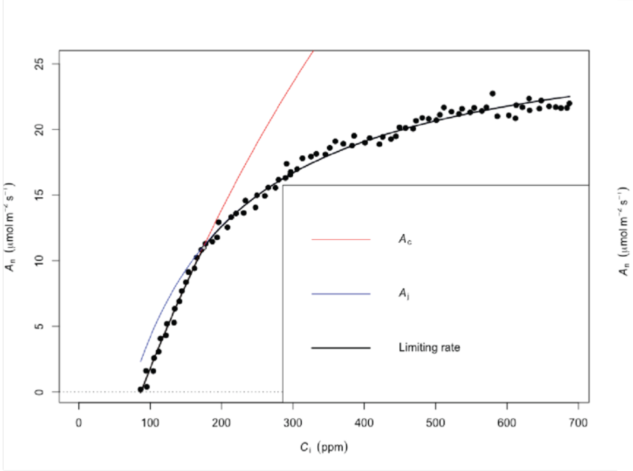


Replicate 4:

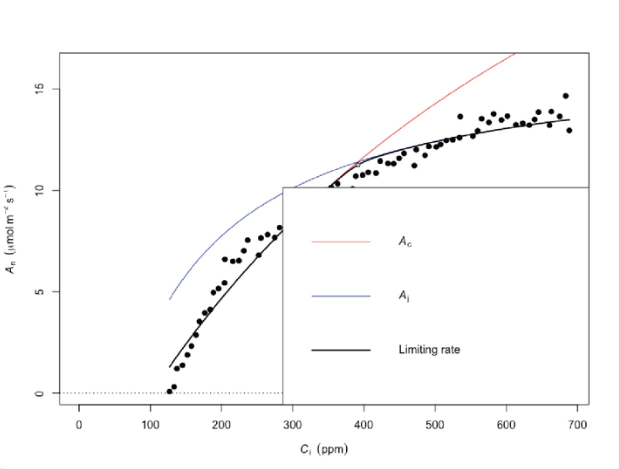

**g**

#### Allele 5

Replicate 1:

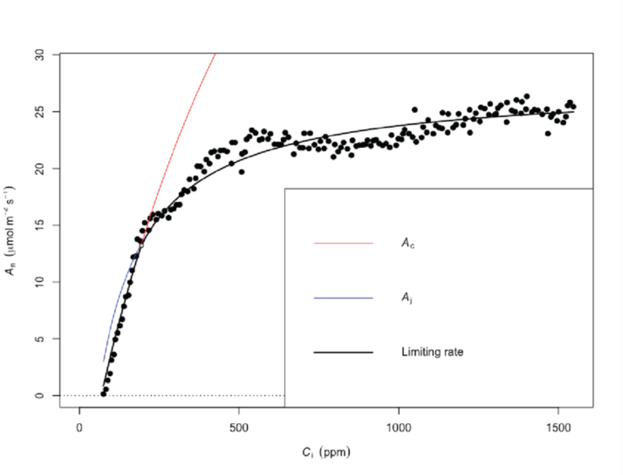


Replicate 2:

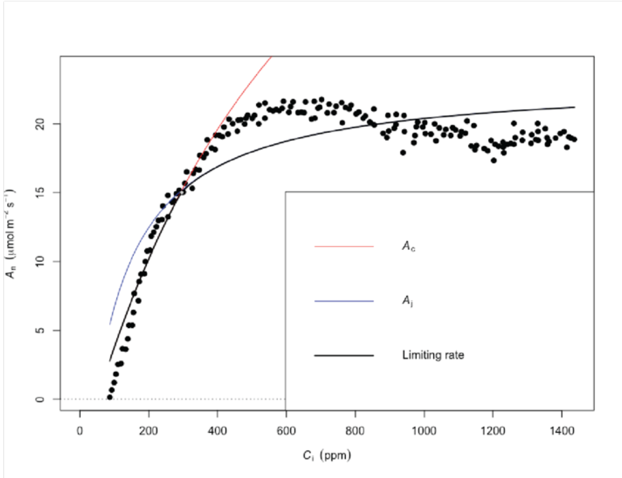


Replicate 3:

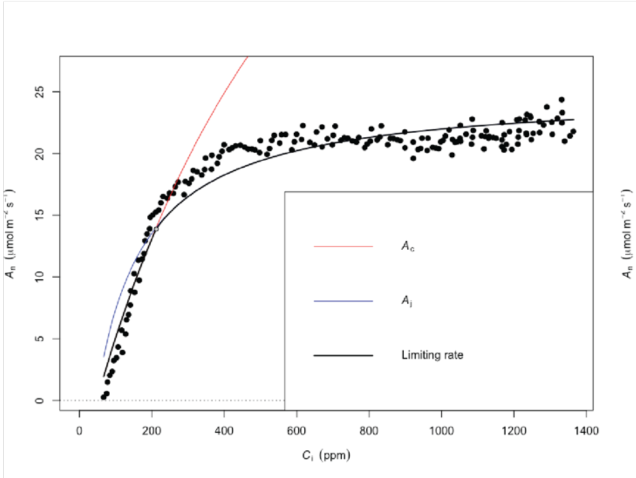


Replicate 4:

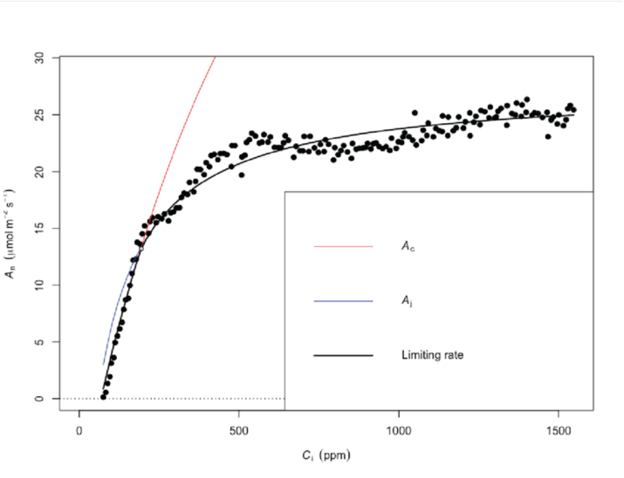

## h

#### Allele 6

Replicate 1:

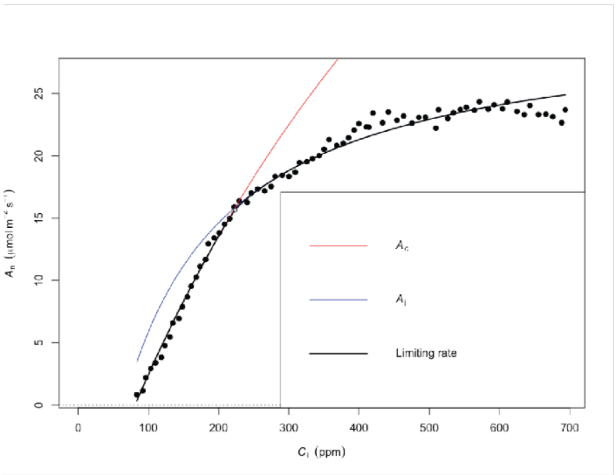


Replicate 2:

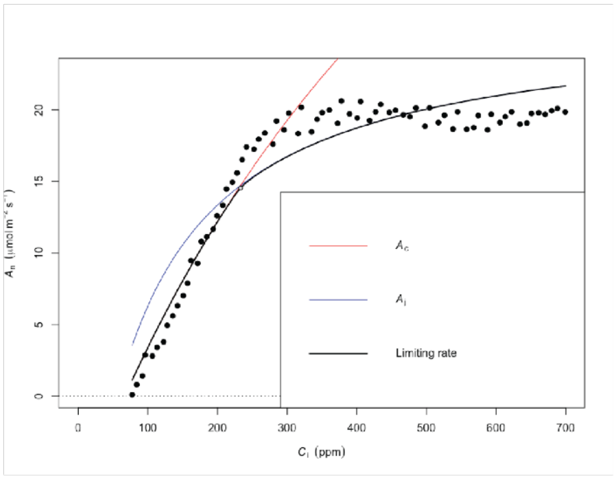


Replicate 3:

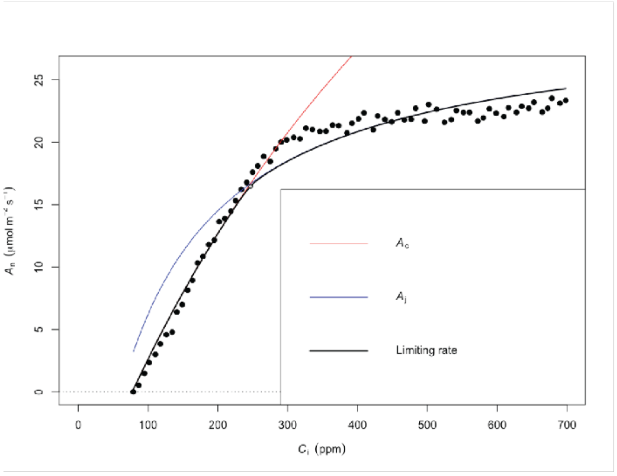


Replicate 4:

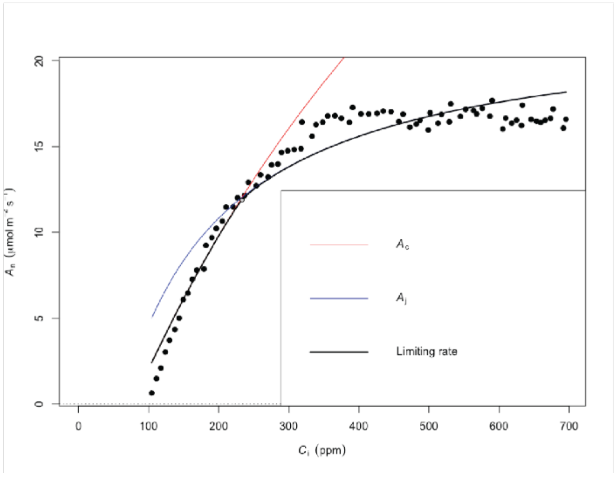


Supplemental Figure 9 | A-Ci curves

The A-Ci curve fit to each biological replicate of (a) *OsSTOMAGEN* (b) Allele 1 (c) Allele 2 (d) Allele 3 (e) Allele 4 (f) Wild type (g) Allele 5 (h) Allele 6. Curves were fit to all Ci <700ppm for which assimilation values >0 using the 'plantecophys' package in Rstudio [35].

Supplemental Figure 10 | Stomatal conductance fluctuating light response curve for each allele

*stomagen*

Replicate 1:

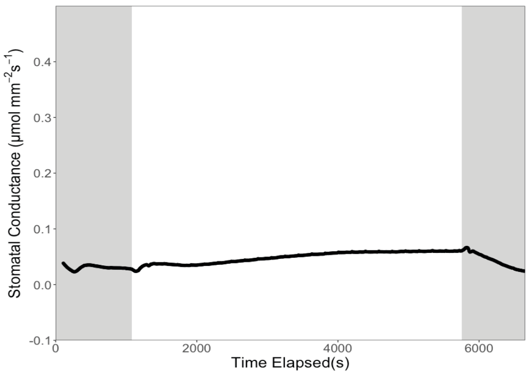


Replicate 2:

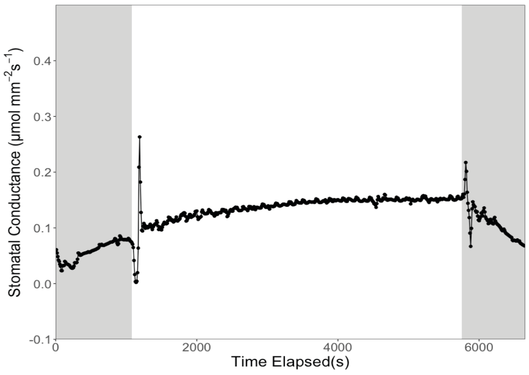


Replicate 3:

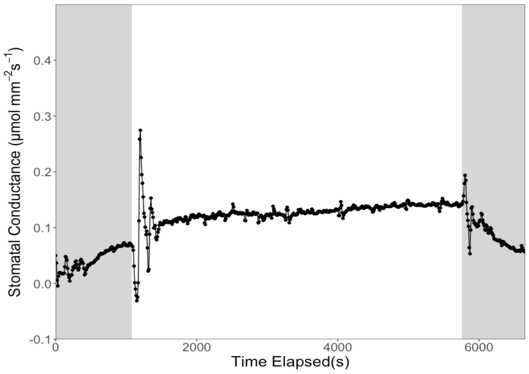


Replicate 4:

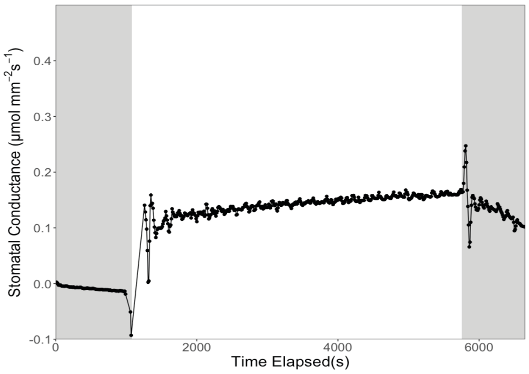

### Allele 1

Replicate 1:

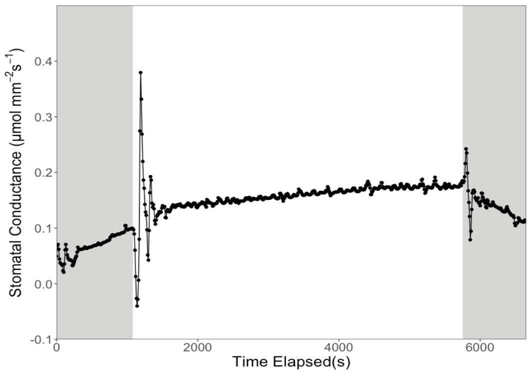


Replicate 2:

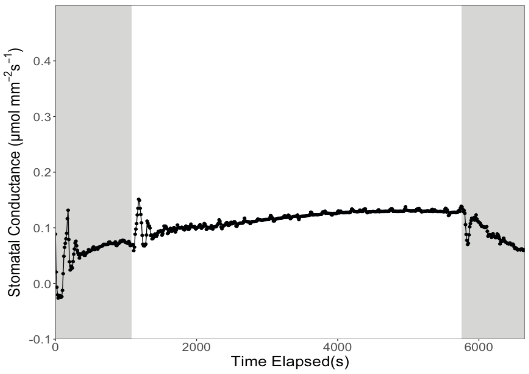


Replicate 3:

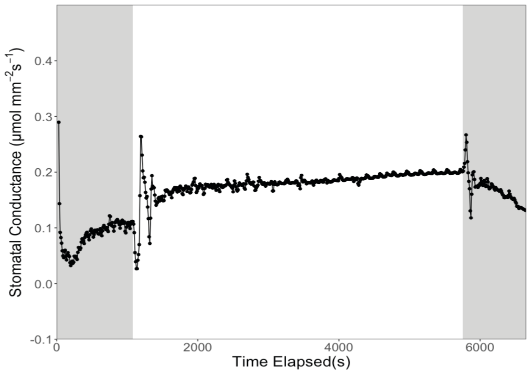


Replicate 4:

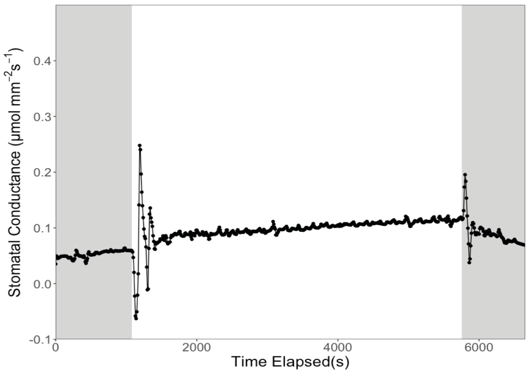

### Allele 2

## UG5

Replicate 1:

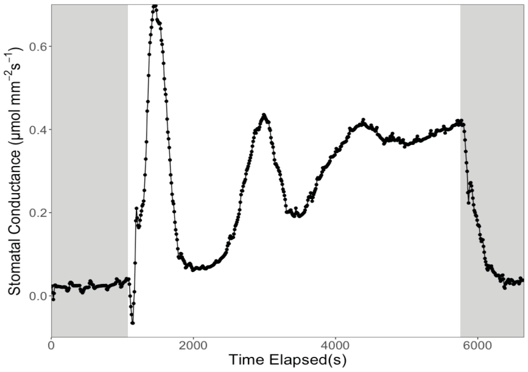


Replicate 2:

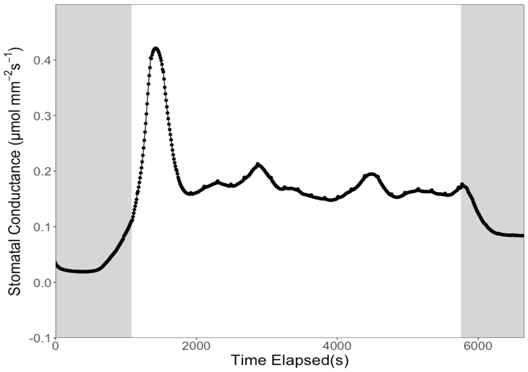


Replicate 3:

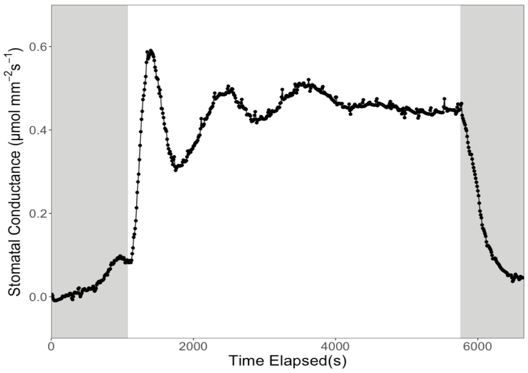


Replicate 4:

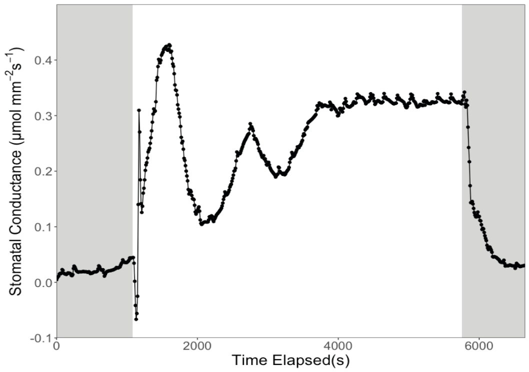

### Allele 3

Replicate 1:

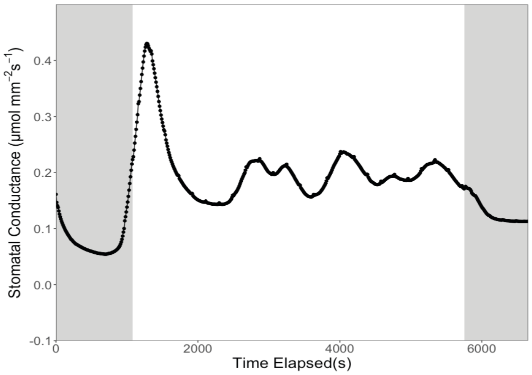


Replicate 2:

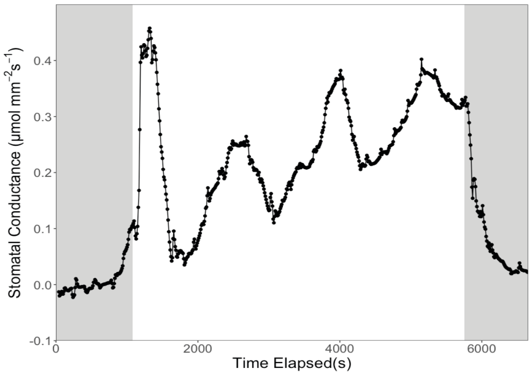


Replicate 3:

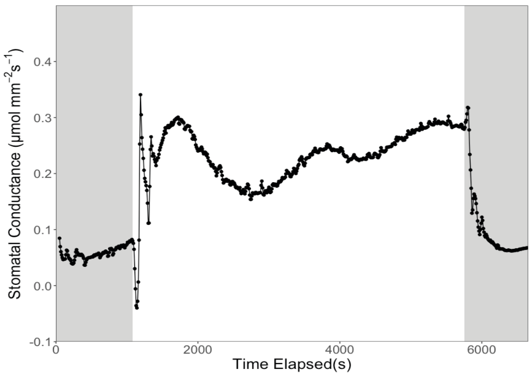


Replicate 4:

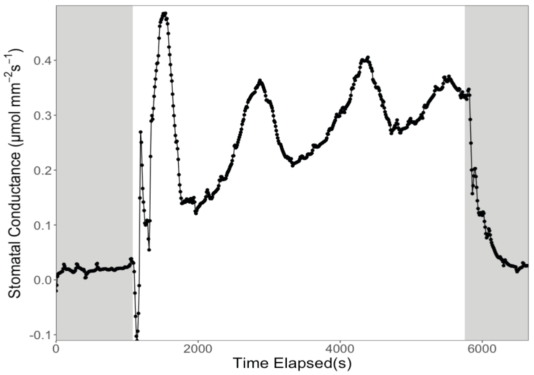

Allele 4

Replicate 1:

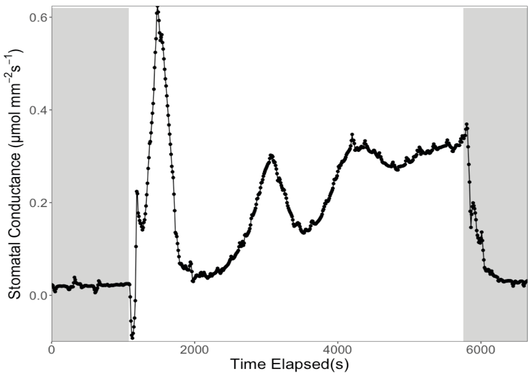


Replicate 2:

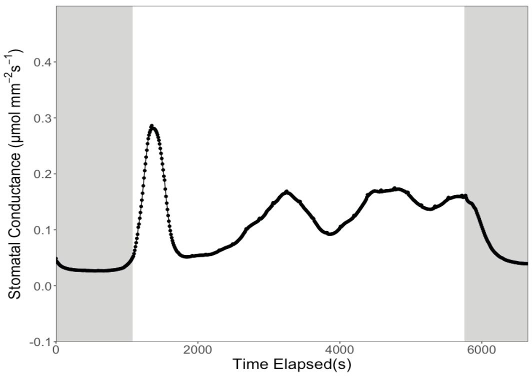


Replicate 3:

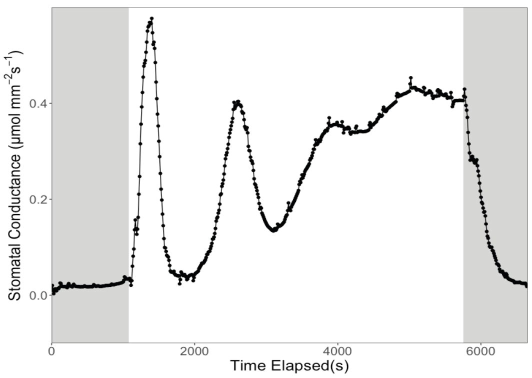


Replicate 4:

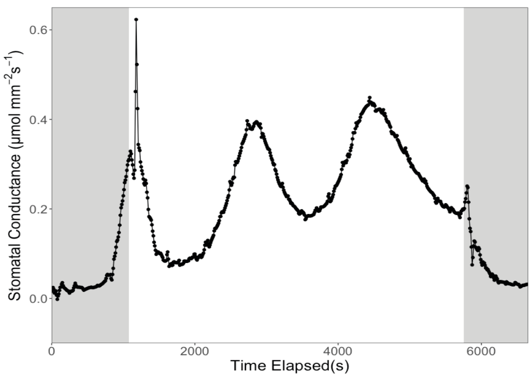

### Wild type

Replicate 1:

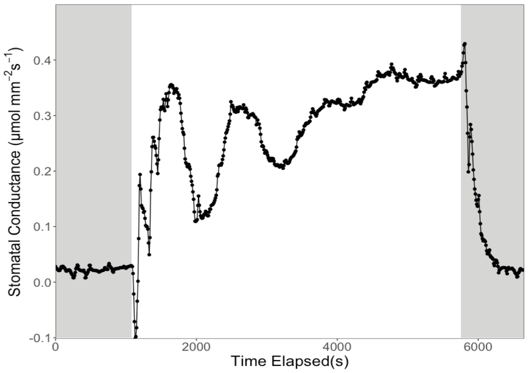


Replicate 2:

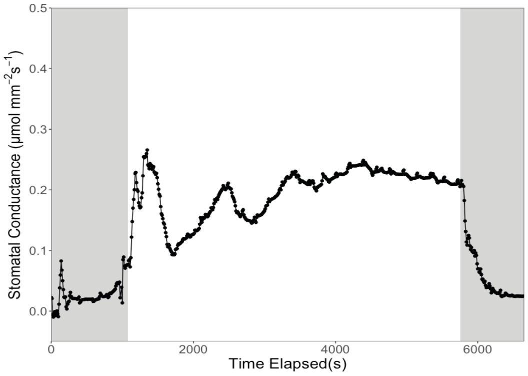


Replicate 3:

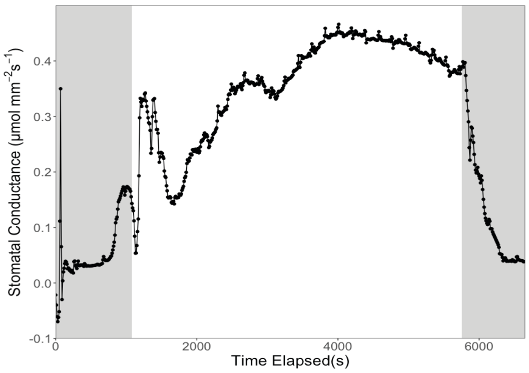


Replicate 4:

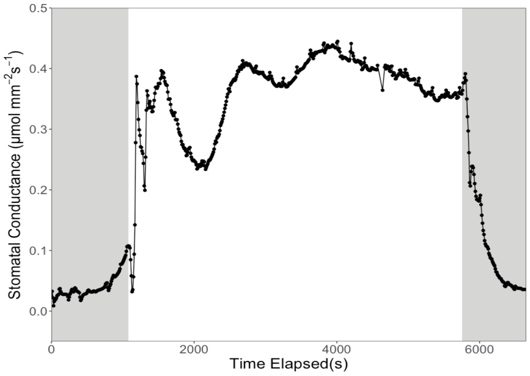

### Allele 5

Replicate 1:

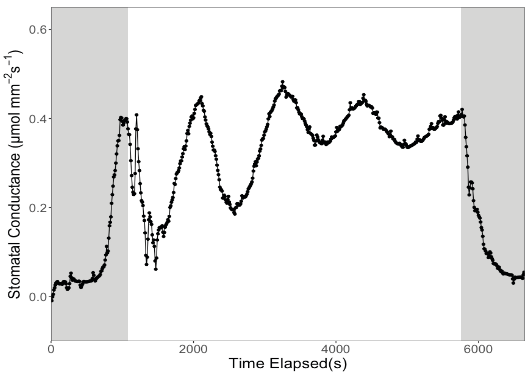


Replicate 2:

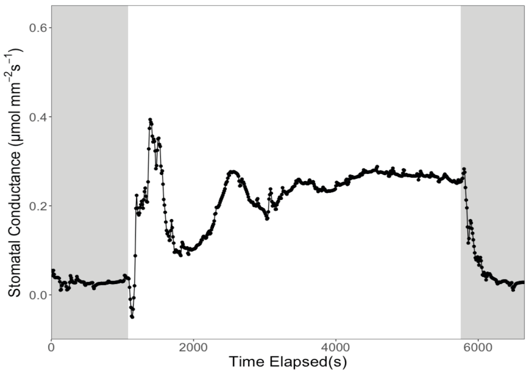


Replicate 3:

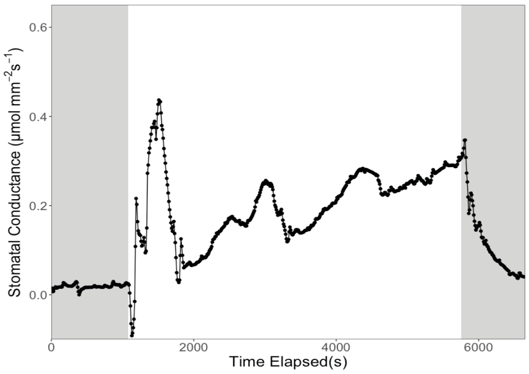


Replicate 4:

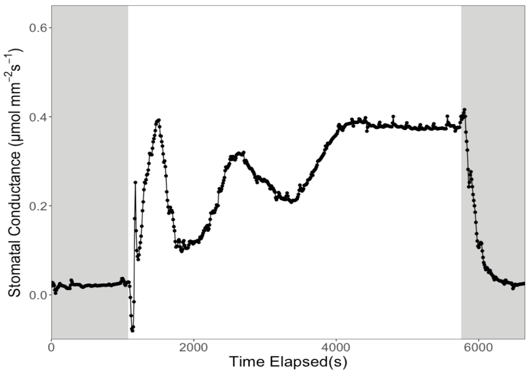

Allele 6

Replicate 1:

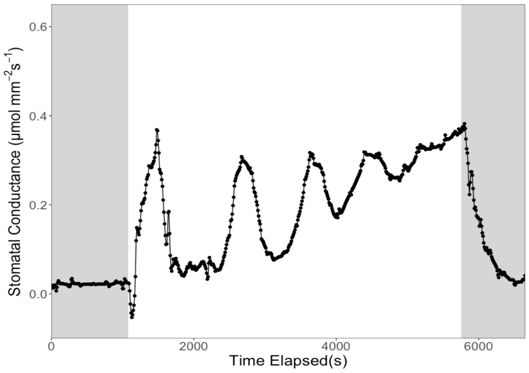


Replicate 2:

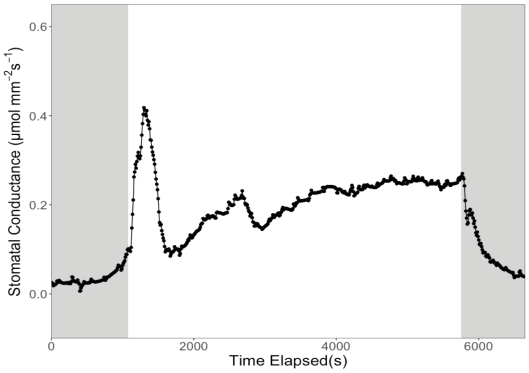


Replicate 3:

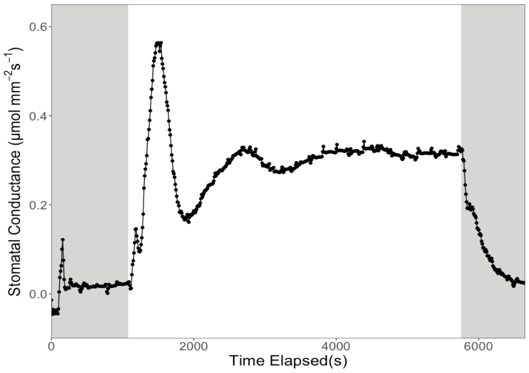


Replicate 4:

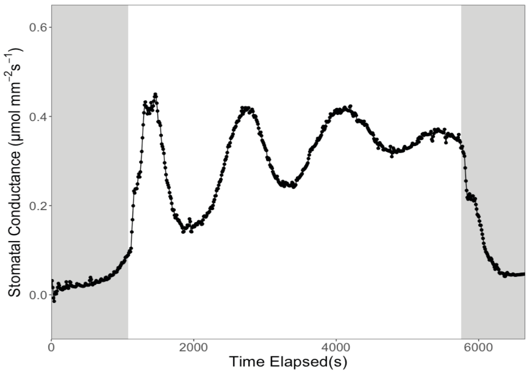

Supplemental Figure 10 | Stomatal conductance response curves in fluctuating light

Stomatal conductance measurements captured over the course of the low-high-low light regime. Data from each of the four biological replicates per genotype are shown. Gray shaded regions indicate low light (100 µmol photons $m^{-2}s^{-1}$) and no-shade regions represent high light of (1500 µmol photons $m^{-2}s^{-1}$).

Table S1: Primer sequences

| Sequence (5' to 3') | Application |
|---|---|
| TAACCTTGAGTTAGATCCAGTGAAGCAAC | Amplifying and subcloning *OsSTOMAGEN* promoter |
| AACCCTTCTTCAAACAAATGGATAGAGAATGG | |
| ATAGTCTCCAGCATTTGCTCCC | Amplifying *OsSTOMAGEN* coding sequence |
| CTGATGCAAAGGGGTACCTGAG | |
| ACCACTTCGACCGCCACTACT | Os*UBQ5* qPCR primers |
| ACGCCTAAGCCTGCTGGTT | |
| TTTCACTCTTGGTGTGAAGCAGAT | Os*eEF-1A* qPCR primers |
| GACTTCCTTCACGATTTCATCGTAA | |
| GCTCGTTGCAATCAAGGGCA | Os*STOMAGEN* qPCR primers |
| GCAGCCTCTCCTTGTTTAGAAC | |

Table S2: Guide sequences arranged from distal through proximal to translation start site

| |
|---|
| TAAAATGTATTTAAAGCTTG |
| TTTGCACGCAATGAAGCATT |
| ATCTTGCAGAAAGCAATTGA |
| ACTTACCGCCTGTTACACGA |
| TGGGCGAGAAAGCAATGAGA |
| GTAGAACAAAAAGAACAAAG |
| GCAGAAGAGCACATGTATAA |
| CAGTGTTGTATAGCGAGAAG |

Table S3: Summary of linear regressions

| Regression of stomatal density by | Correlation Coefficient | p-value | Equation of the line of best fit |
|---|---|---|---|
| Guard cell length | -0.56 | 0.15 | y=24-0.0087x |
| Carbon assimilation | 0.92 | 0.001 | y=10+0.027x |
| Stomatal Conductance | 0.98 | 0.000009 | y=0.085+0.00062x |
| Intrinsic water-use efficiency | -0.95 | 0.00023 | y=95-0.079x |
| ΦPSII | 0.84 | 0.0087 | y=0.18+0.00017x |
